# Distinct brain networks for remote episodic memory depending on content and emotional value

**DOI:** 10.1101/2022.09.16.508241

**Authors:** Anne Auguste, Nicolas Fourcaud-Trocmé, David Meunier, Alexandra Gros, Samuel Garcia, Belkacem Messaoudi, Marc Thevenet, Nadine Ravel, Alexandra Veyrac

## Abstract

The mechanisms that underlie the storage of old nontraumatic episodic memories remain enigmatic because of the difficulty in modelling this particular type of memory in humans and animals. Episodic memories combine incidental and occasional “What-Where-When/In which context” multisensory information and can be stored for long periods of time. Here, using a task in rodents that models human episodic memory, including odour/place/context components, we applied advanced behavioural and computational analyses and brain imaging of *c-Fos* and *Zif268* to characterize remote episodic memories and their engrams for the first time. We show that the content and accuracy of memories vary across individuals and depend on the emotional relationship with odours experienced during episodes. Activated brain networks reflect the nature and content of remote episodic memories and their transformation over time, and emotional cortico-hippocampal networks play critical roles in maintaining vivid memories.

## Introduction

The mechanisms underlying the storage of old nontraumatic episodic memories remain poorly understood. Episodic memories are formed incidentally by associating several pieces of information (What? Where? When/In which context?)^1–3^ and may recruit large brain networks to form engrams that evolve over time. Complex because highly detailed, malleable because formed without extensive learning, but robust to face the passage of time^1, 2^, episodic memories defy our knowledge of their supportive engram and fate. Understanding why and how episodic memories, which are critical for individuals, are maintained and transformed over time is an important issue for understanding normal and pathological brain function^4^.

The characteristics and brain networks associated with human episodic memories have been studied in healthy subjects and patients by using paradigms with autobiographic or laboratory approaches^5–7^. However, the results differ depending on the approach used^8^, and many questions remain unexplored or highly debated, especially those concerning the evolution of episodic memory engrams over time. Two main theoretical frameworks are still evolving and offer contrasting views on the role and interactions of the medial temporal lobe and cortical areas in the consolidation of remote declarative memories^9–12^.

Studies on animal models of episodic memory have provided crucial information about the general characteristics of episodic memory^4, 13^; however, these studies used experimental paradigms that neglect some important features of human episodic memory. Some studies have shown that rodents can encode events without training^14–16^ and form complex episodic *What-Where-When*^17–20^ or *What-Where-In Which Context* representations^3, 21–24^. However, these paradigms require extensive training during encoding and/or fail to test memory durability over time. While some studies have explored the fate of remote memories, they did not associate the three components specific to episodic memory^25–29^. Consequently, brain networks associated with remote episodic memories are *Terra incognita* in animals, and this gap should be investigated to explore mechanistic aspects that are difficult to address in humans. Thus, we developed a new episodic memory task for rats that models human episodic memory and allows old memories to be investigated^30, 31^.

In the present study, rats form remote episodic memories by experiencing immersive life episodes with olfactory, spatial and contextual information. We applied advanced behavioural and computational analyses and observed that despite high sensitivity to interference during recall, remote episodic memory remains robust over time. Similar to humans, we found that the information content in episodic memories varies across individuals. Reproducible memory profiles correlate with the initial emotional experience with odours during episodes. By using cellular imaging of *c-Fos* and *Zif268* in various brain regions and functional connectivity analyses, we provide the first evidence that remote episodic memory networks reflect the information content and level of detail included in episodic memories. We show that the more complete the recollection, the larger the network recruited during recall, with a set of brain areas that represent the experienced information and emotional brain networks related to odours that are critical for maintaining an accurate remote memory. The remote episodic memory engram remains highly dynamic since synaptic plasticity processes are induced during recall, thus supporting post-recall memory updates and reinforcement.

## Results

### Paradigm for exploring remote episodic memory in rats

After adapting to the EpisodiCage (Fig. 1a-b) and humdrum sessions (*Routine*; Fig. 1c), rats experienced two episodes in different multisensory environments^30, 31^ (Fig. 1d). During the episodes, which lasted a maximum of 40 min, rats freely visited two ports that delivered specific odours associated with pleasant or unpleasant drinking solutions. Different odours and active ports are associated during each episode (Episode [E1]: odours O1/O2 with ports P2/P3; Episode [E2]: odours O3/O4 with ports P1/P4). Thus, in a given context (“*In which context*” information), the rats incidentally encode which port (“*Where*” information) and which odour (“*What*” information) is associated with a pleasant or unpleasant experience. To assess remote episodic memory, a recall test session was conducted 30 days after the episodes by placing rats in the same situation as in Episode 2 (Fig. 1e).

**Figure 1.**
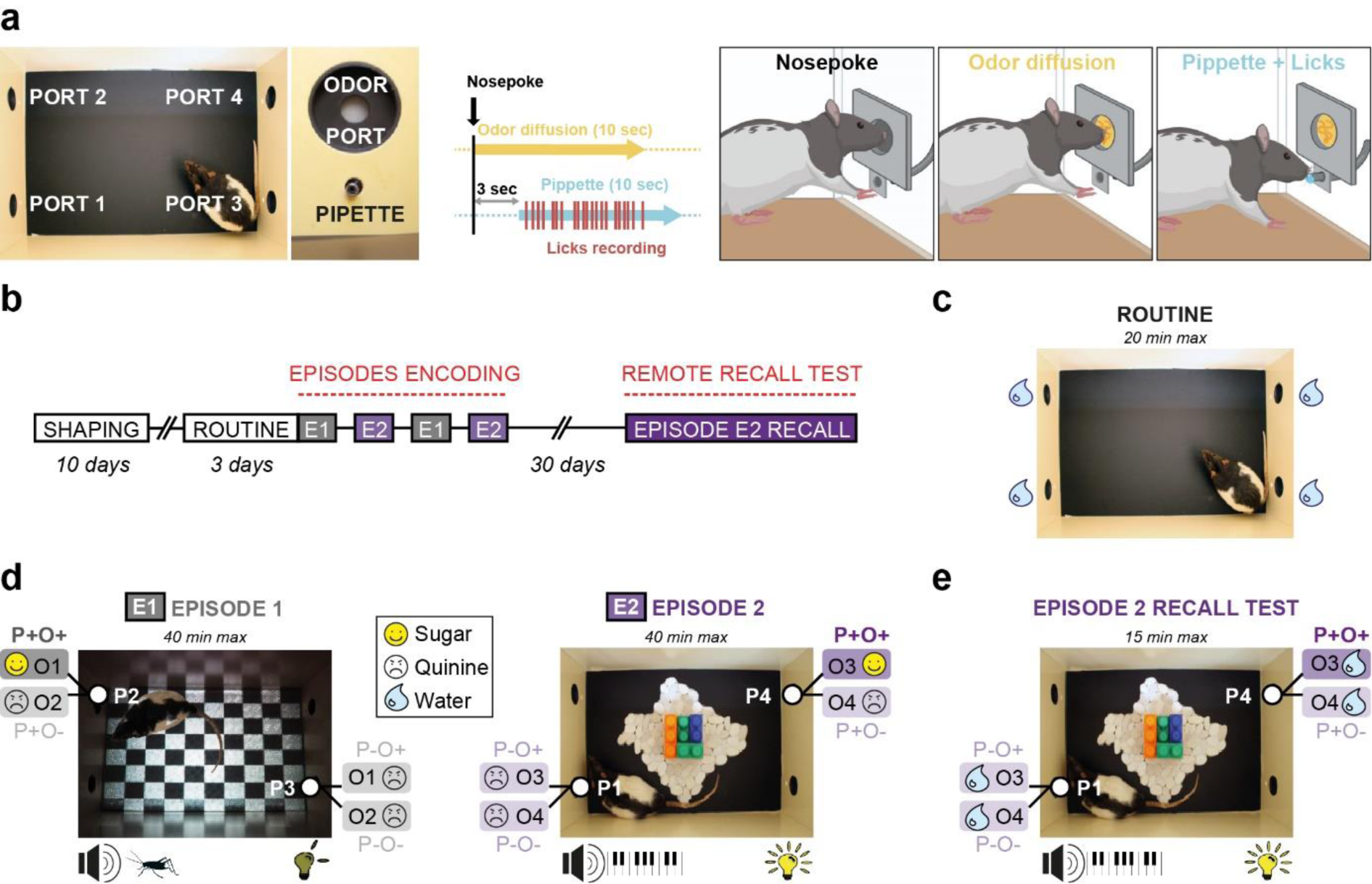
Experimental design for exploring remote episodic memory in rats. **(a)** The EpisodiCage is equipped with 4 odour ports, and each port is associated with a motorized pipette. During a visit to a port, nose pokes are detected and induce 10 sec of odour diffusion. The pipette is introduced into the cage below the port 3 sec after the nose poke, and a drinking solution is delivered for 10 sec. All licks performed by the rat are recorded. **(b)** Timeline of the task: after a 10-day *Shaping* phase to the functioning of the EpisodiCage and 3 daily *Routine* sessions, the rats are exposed twice to two distinct episodes: Episode 1 (E1) and Episode 2 (E2). Their remote episodic memory was tested 30 days after the episodes during a recall session in the context of E2. **(c)** During routine sessions (20 min maximum per day), rats are placed in a neutral version of the environment without odour, and only water is dispersed through the 4 ports. **(d)** During the episodes, which last a maximum of 40 min per session, rats form episodic memories by experiencing two distinct episodes characterized by their environmental context (*Which context*), the location of the active ports (*Where*) and the odours diffused by the ports (*What*). Episode 1: dark environment with sounds of crickets at night, P2 and P3 are active and diffuse odours O1 or O2. Episode 2: bright environment, Legos and pebbles on the floor, piano music, P1 and P4 are active and deliver odours O3 or O4. During each episode, the rats are presented with a pleasant sugar solution at only one port, paired with one odour (yellow smiley; P+O+ configuration with O1 on P2 for E1 and O3 on P4 for E2), while they are presented with an unpleasant bitter solution in all other configurations (black smiley; P+O-; P-O+; P-O- with O2 on P2 and O1/O2 on P3 for E1; O4 on P4 and O3/O4 on P1 for E2). **(e)** Remote episodic memory is tested 30 days after episodes by placing the rats in the context of E2 for a limited period (15 min maximum). During the recall test, only water is delivered to evaluate what the rats remembered.

### Rats form and recollect a robust remote episodic memory

During each episode, rats preferentially visited the good port (P+) (Fig. 2a; *P*_Wilcoxon_<0.001 visits P+ *vs.* P**-** for all episodes) and licked the P+O+ configuration, which was the only configuration that provided sugar solution, more often (Fig. 2b; *P*_Friedman_<0.005, *P*_Wilcoxon_<0.005 licks P+O+ *vs.* other configurations). Furthermore, individual experiences during each episode appeared roughly homogeneous (Fig. 2c).

**Figure 2.**
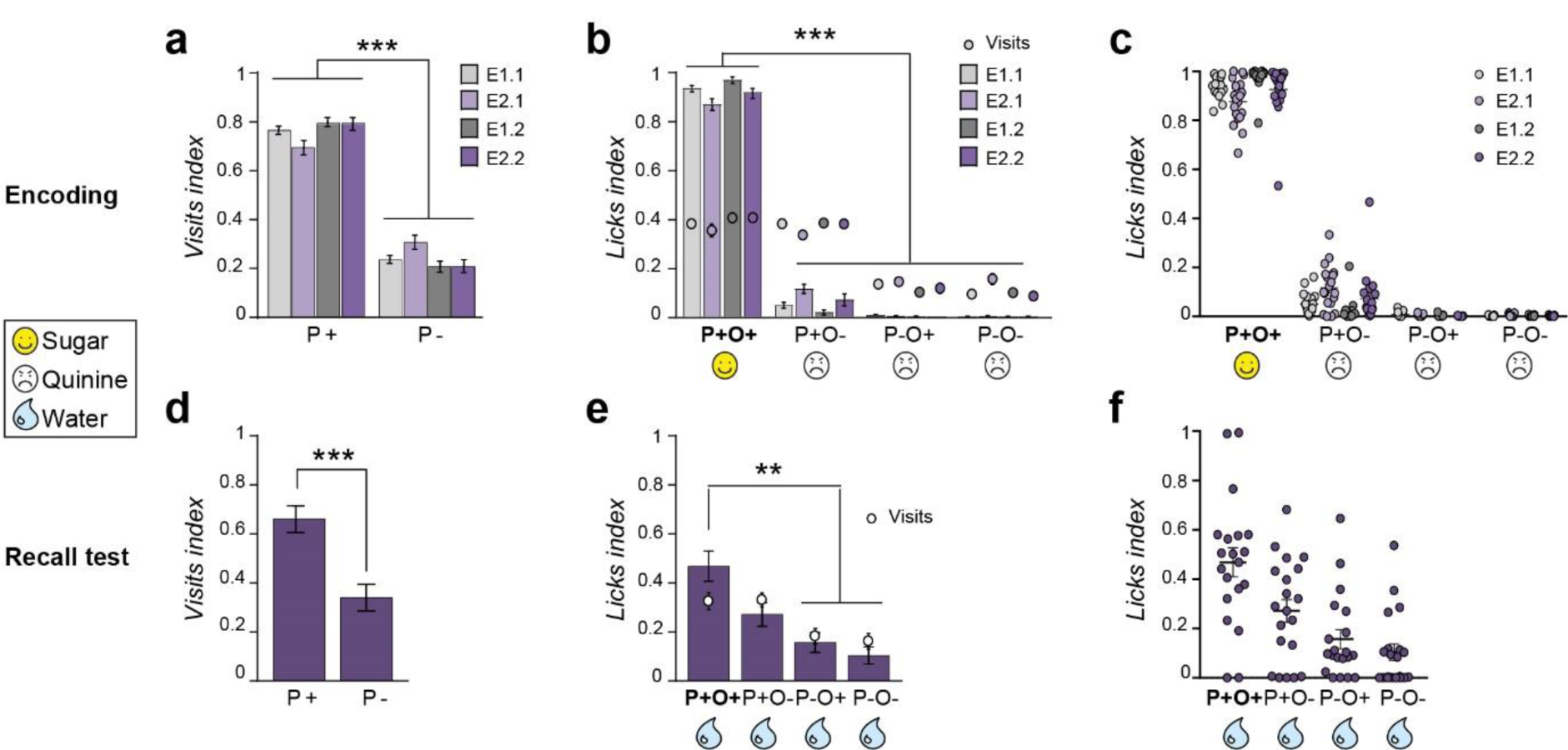
Rats form and recollect a robust remote episodic memory. **(a)** Index of visits to each port during episode encoding. Rats explored the good port P+ significantly more often than the other ports during all episode encoding sessions. **(b)** Indices of visits and licks for each configuration during encoding. Rats licked the P+O+ configuration (good odour on the good port), which was associated with a pleasant experience of sugar solution, more than the other configurations, which were associated with an unpleasant experience of bitter solution (P+O-: good port, bad odour; P-O+: bad port, good odour; P-O-: bad port and bad odour). **(c)** Individual data of the licks index during encoding were relatively homogeneous. **(d)** Index of visits to each port during the recall test at 30 days, showing that the rats remembered the good port P+. **(e)** Indices of visits and licks for each configuration during the recall test. Rats remembered the good configuration P+O+, which was associated with sugar during E2. **(f)** Individual data of the licks index for each configuration during the recall test showed high individual variability in the remote episodic memory content. Group data are expressed as the mean ± SEM (n = 19). ***p<0.005, **p<0.01, Friedman tests followed by Wilcoxon tests for statistical comparisons between configurations. E1.1 (first episode E1); E2.1 (first episode E2); E1.2 (second episode E1); E2.2 (second episode E2).

To examine remote memory of previous episodes, rats were placed in the Episode 2 context 30 days later with only water delivered (Fig. 1e). The rats’ performance represents what the rats expected to drink and thus their episodic memory (Fig. 2d-f). The group performance results showed that the rats remembered the episodic association since they explored the good port P+ significantly more than the other port (Fig. 2d; *P*_Wilcoxon_<0.001 visits P+ *vs.* P-) and licked the good configuration P+O+, which was previously associated with sugar, more often (Fig. 2e; *P*_Friedman_=0.005, *P*_Wilcoxon_<0.01 licks P+O+ *vs.* P**-**O+ and P**-**O**-**, *P*_Wilcoxon_=0.056 licks P+O+ *vs.* P+O**-**). Interestingly, in contrast to the encoding results (Fig. 2c), individual variability in the remote episodic memory performance was more pronounced (Fig. 2f).

### The diverse content of remote episodic memory

In contrast to recent memories, for which 100% of the rats recollected all episodic information^30^, three distinct memory profiles were observed 30 days after the episodes (Fig. 3a).

**Figure 3.**
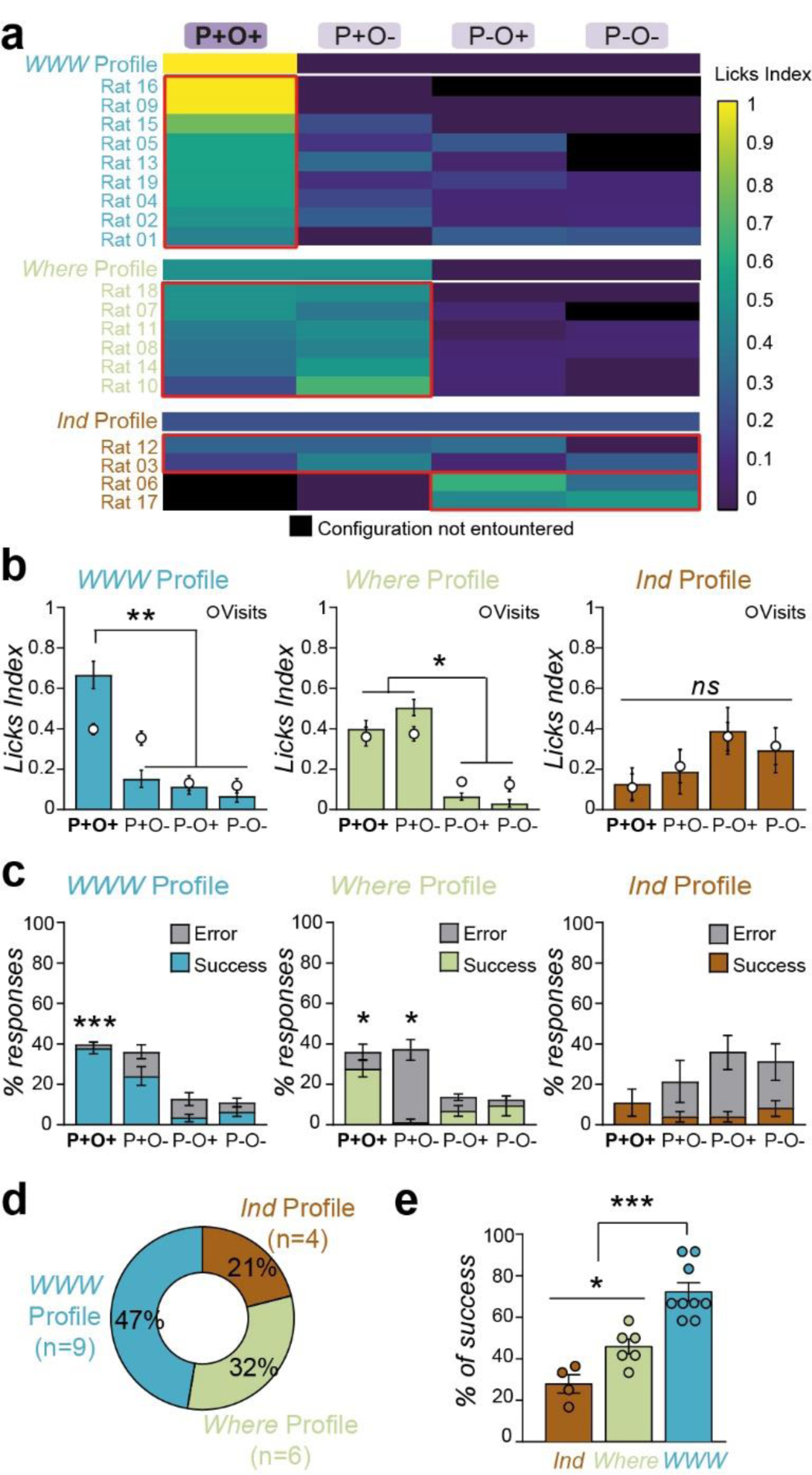
The different contents of remote episodic memory. **(a)** Colour matrix of the individual licks index, showing the three memory profiles observed during the recall test 30 days after episode encoding. Rats with *What-Where-Which Context* (*WWW*) profile remembered the good P+O+ configuration in the episode. Rats with a *Where* memory profile remembered only the correct location (P+O+ and P+O- configurations). Rats with an *Indeterminate* memory profile (*Ind*) explored and drank from all configurations. **(b)** Mean of licks and visits indices and **(c)** percentages of successes and errors for the different memory profiles. *WWW* rats licked the P+O+ configuration significantly more and made more correct responses on the P+O+ configuration. *Where* rats visited and licked the P+ configuration significantly more often, regardless of what the odour was, and made more errors on P+O-. *Indeterminate (Ind)* rats did not visit or lick more or show clearer responses on any configuration. **(d)** Proportions of different memory profiles. **(e)** Overall success rate during the recall test for each memory profile. Group data are expressed as the mean ± SEM (n*_WWW_* = 9, n*_Where_* = 6, n*_Ind_* = 4). *p<0.05; **p<0.01; ***p<0.005, Friedman tests followed by Wilcoxon tests for statistical comparisons between configurations or types of responses and Kruskal‒ Wallis test followed by Mann‒Whitney U tests for statistical comparisons between memory profiles. P+O+ (good port and good odour); P+O- (good port and bad odour); P-O+ (bad port and good odour); P-O- (bad port and bad odour).

Despite their limited experience, half of the rats (47%) recollected all the episodic information, presenting “*What**-**Where**-**Which context*” (*WWW*) profile during the test (Fig. 3a, d). *WWW* rats licked the P+O+ configuration significantly more (Fig 3b; *P*_Friedman_<0.001, *P*_Wilcoxon_=0.008 licks P+O+ *vs.* other configurations) and made more correct responses (Fig 3c; *P*_Wilcoxon_=0.001 success *vs.* error on P+O+). One third of the rats (32%) remembered only the spatial information (*Where* rats, Fig 3a, d), visiting and licking P+ significantly more, regardless of whether the odour was O+ or O**-** (Fig. 3b; *P*_Friedman_=0.002, *P*_Wilcoxon_=0.028 licks on P+O+ and P+O**-** *vs.* P**-**O+ and P**-**O**-**, *P*_Wilcoxon_>0.05 licks P+O+ *vs.* P+O**-**). *Where* rats also made more correct responses on P+O+ and more errors on P+O- (Fig. 3c; % success *vs.* error *P*_Wilcoxon_=0.046 on P+O+, *P*_Wilcoxon_=0.027 on P+O**-**). Finally, 21% of the rats showed an *Indeterminate (Ind)* memory profile (Fig. 3a, d), as they did not visit or lick any port significantly more often (Fig. 3b; *P*_Friedman_=0.601) and showed no clear responses to any given configuration (Fig. 3c; *P*_Friedman_=0.12). Thus, the precision of remote episodic memory shows a significant increase across memory profiles (Fig. 3e; *P*_Kruskall-wallis_=0.001, *P*_Mann-Whitney_=0.024 *Indeterminate vs. Where*, *P*_Mann-Whitney_=0.002 *WWW vs. Where* and *Indeterminate*). These specific memory profiles remained highly robust throughout the recall test session (Supplementary Fig. 1) and are not related to biased experiences during episode encoding since rats with all three memory recall profiles encoded episodic associations (Supplementary Fig. 2).

These results are the first to show that, after very long delays, rats can remember time-limited life events according to three categories of multisensory information (odour/place/context), although the individual variability during recall suggests that remote episodic memory is a system with diverse content.

### Despite time and interference during recall, certain remote episodic memory components are preserved

In daily life, recall of past events generally occurs in contexts different from those of the initial encoding, leading to implicit comparisons and information sorting according to distinct but similar life episodes. One example is when humans return to a given place with different people for another occasion and attempt to recollect information about the initial event. Here, we examined whether remote episodic memory in rats can address this challenging situation and how different memory components are preserved over time.

A new group of rats was exposed to the previously described episodes and tested 30 days later in a challenging recall situation by using 4**-**Port test (Fig. 4a). In this situation, rats were returned to the context of Episode 2, including its odours and ports (*In context* configurations, *IC*); however, the odours and ports of Episode 1 were also present (*Out of context* configurations, *OC*)^30^. Episodic encoding occurred in a similar manner as previously described (Fig. 4b; *P*_Friedman_<0.001, *P*_Wilcoxon_=0.005 licks P+O+ *vs.* other configurations; *P*_Mann-Whitney_>0.05 for encoding data between the two groups of rats, Figs. 2b and 4b). However, during the recall test, individual memory performance varied more in this situation than when the test occurred in the same context as Episode 2 (Fig. 4c; *P*_Friedman_=0.039, *P*_Wilcoxon_>0.05 between configurations); 70% of the rats recollected at least the contextual information of Episode 2 (*WWW/Where*/*Indeterminate*-*In context* profiles), 20% became confused due to the contextual elements of both encoding episodes (*Where*/*Indeterminate***-***Out of context* profiles) and 10% displayed indeterminate profile (Fig. 4 d, e). Among the rats that handled the context interference, 20% remembered the whole episode (*WWW***-***In context* profile), 20% retrieved only the place information (*Where***-***In context* profile) and 30% recollected only the context information (*Ind***-***In context* profile).

**Figure 4.**
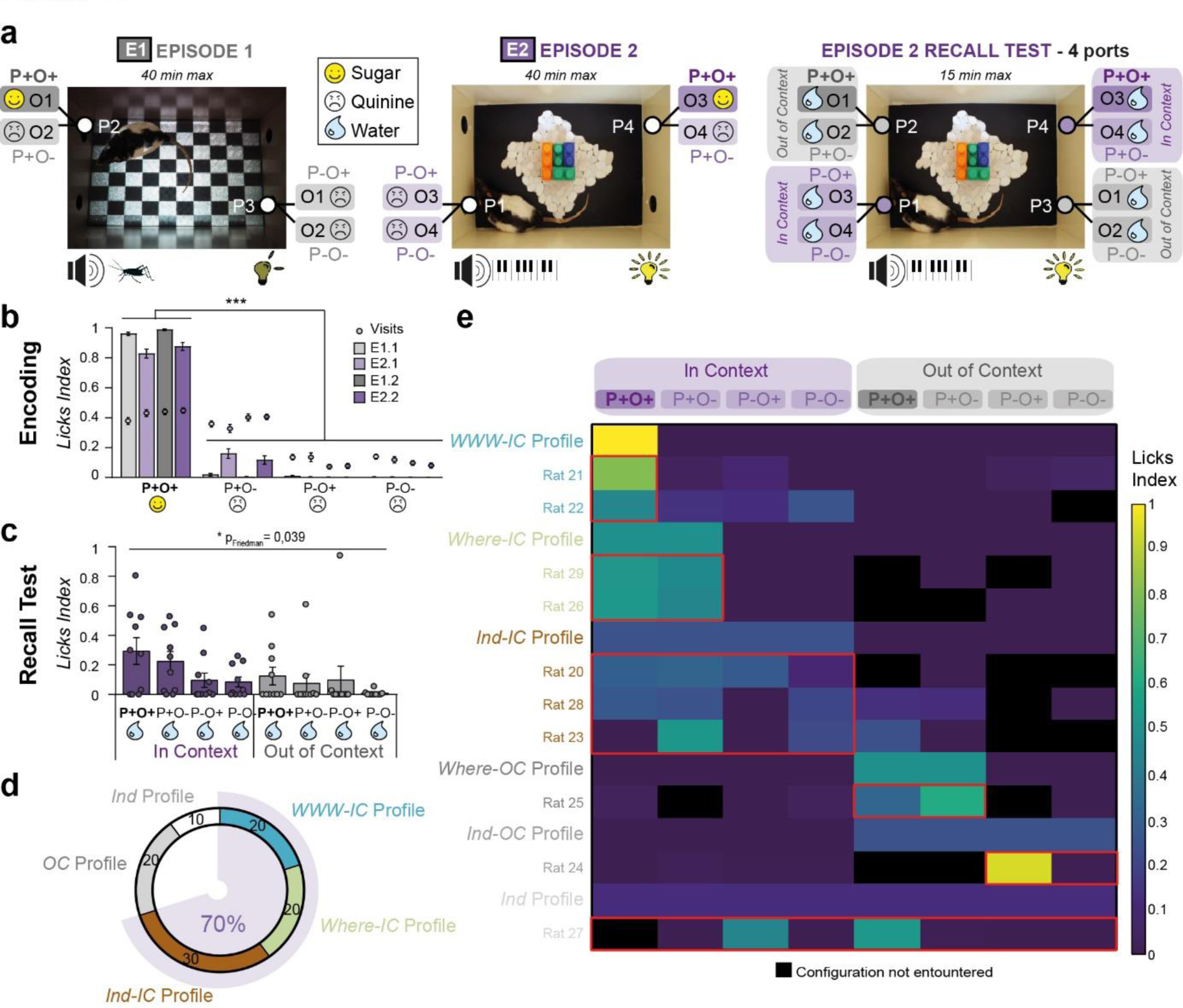
Remote episodic recollection is sensitive to interference, but certain memory components are preserved. **(a)** After similar encoding sessions for E1 and E2, a new group of rats (n = 10) was subjected to a more challenging version of the recall test that used 4 ports. Rats were placed in the E2 context with its odours and ports (*IC, In context* configurations: P+O+, P+O-, P-O+, P-O-); however, the environment also included the odours and ports of E1 (*OC*, *Out of Context* configurations: P+O+, P+O-, P-O+, P-O-). **(b)** Licks and visits indices during episode encoding. Rats licked the P+O+ configuration, which was associated with a pleasant experience of sugar solution, more than the other configurations, which were associated with an unpleasant experience of bitter solution. **(c)** Group and individual data of the lick index during the 4-port recall test. The group data did not reveal significant differences between configurations, and the individual data were more variable in this test situation. **(d, e)** Proportions of the different memory profiles during the 4-port recall test. Colour matrix of the individual licks index on the 8 configurations. In this situation, 70% of the rats recollected at least the contextual information of E2 (all *IC* profiles), 20% of the rats were confused with the context (all *OC profile*) and 10% of the rats did not appear to remember any relevant information (*Ind* profile). Group data are represented as the mean ± SEM. *p<0.05, ***p<0.005, Friedman tests followed by Wilcoxon tests for statistical comparisons between configurations. *Ind* (*Indeterminate*); *IC* (*In Context*); *OC* (*Out of Context*); P+O+ (good port and good odour); P+O- (good port and bad odour); P- O+ (bad port and good odour); P-O- (bad port and bad odour).

Our results show that despite its robustness across time, remote episodic memory is highly sensitive to retrieval interference (20% of rats were *WWW* in the 4-Port test, Fig. 4d, e *versus* 47% in the similar recall situation, Fig. 3d). Moreover, context and spatial information are preserved more over time. In the 4-Port test, 70% of the rats remembered *In context* information after 30 days, similar to the 80% of rats that remembered in 1**-**day recall test^30^; 20% remembered only the context and spatial information after 1 and 30 days; and 20% of the rats were *WWW* at 30 days *versus* 50% at 1 day.

### Recollection of a complete remote episodic memory is related to an unpleasant experience during the first episode

Since we observed several memory recall profiles with highly reproducible proportions in different experiments, we investigated their potential origin. Individual variability in episodic memory content could emerge from *i)* distinct memory retrieval abilities during the test, *ii)* distinct memory consolidation processes during the 30-day retention, or *iii)* different experiences during episode encoding. In the present study, we explored the potential contribution of individual experiences during encoding.

We performed a computational study to analyse whether the behavioural data collected during encoding could be used to predict remote episodic memory performance. We applied linear models and determined the best sets of two or three encoding parameters for describing remote memory performance (% of successes during the test). The results are displayed in Table 1, showing that the only explanatory variable in all selected models was the parameter corresponding to avoiding drinking from port P+ in the presence of odour O- during the first presentation of Episode 1 (index of correct rejection on P+O- during E1.1). This parameter was negatively correlated with success during recall, which reflects memory precision (*P*_Spearman_=0.0024; Rho=-0.632) (Fig. 5a), suggesting that during the first exposure in Episode 1, the more rats experienced the unpleasant drinking solution in the good port, the better their remote episodic memory.

**Figure 5.**
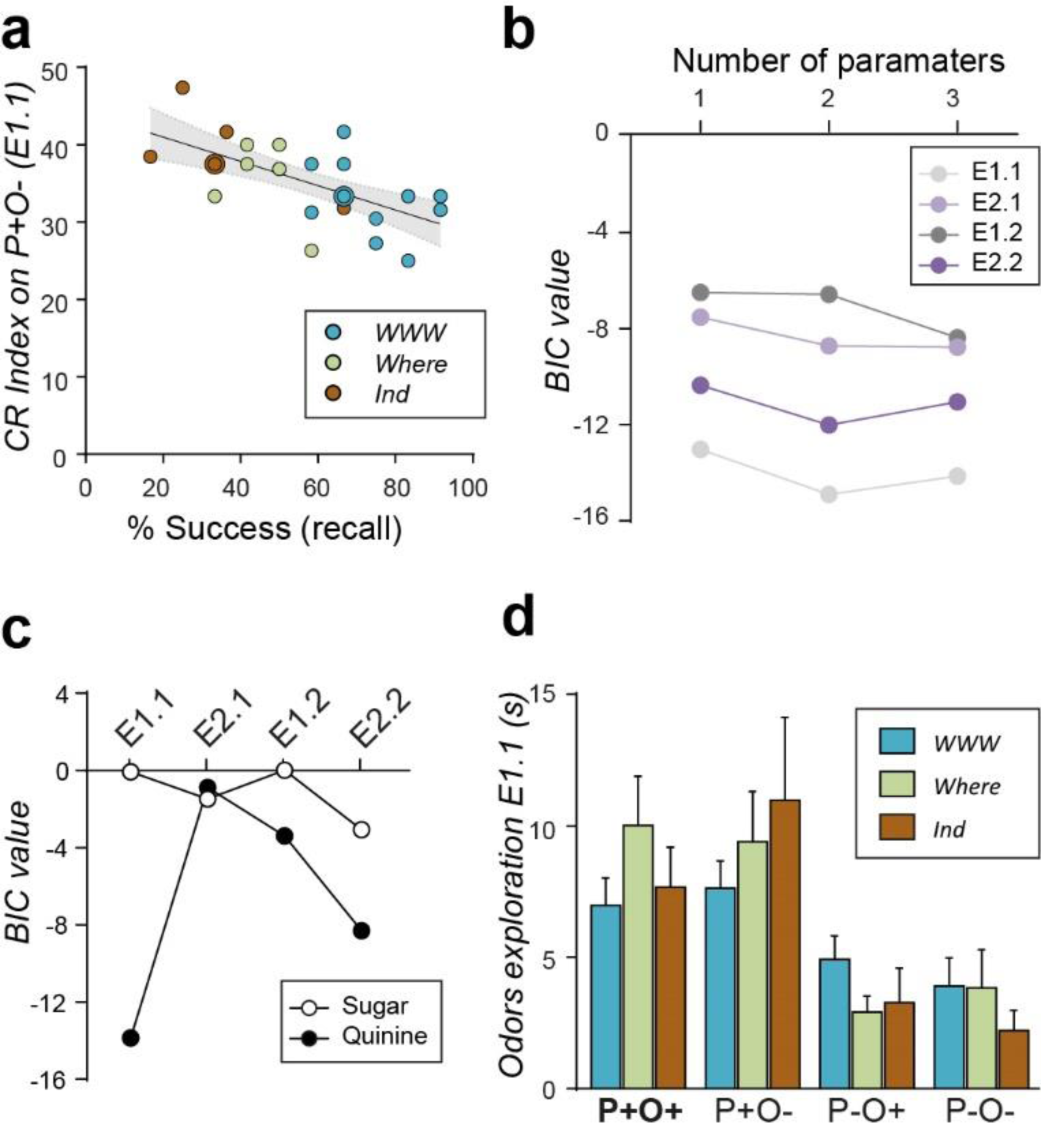
Accuracy of remote episodic memory depends on negative experience during the first episode. **(a)** Remote episodic memory performance (success rate during the recall test) was negatively correlated with the correct rejection index on P+O- during E1.1 (n = 24; p_Spearman_=0.0024; Rho=- 0.63). **(b)** BIC value of each regression model (for all episode sessions), showing the critical role of the initial episode E1 (E1.1) in the remote memory performance, with lower BIC values regardless of the number of parameters used in the model. **(c)** BIC value of each regression model for sugar *versus* quinine experience during each episode session, showing that an unpleasant experience during the first episode E1 (E1.1) was a strong predictor of remote episodic memory precision. **(d)** The duration of odour exploration for each configuration during the first episode E1 was similar for all memory profiles. Group data are represented as the mean ± SEM (n*_WWW_* = 12, n*_Where_* = 7, n*_Ind_* = 5). Kruskal‒Wallis tests for statistical comparisons between memory profiles. E1.1 (first episode E1); E2.1 (first episode E2); E1.2 (second episode E1); E2.2 (second episode E2); P+O+ (good port and good odour); P+O- (good port and bad odour); P-O+ (bad port and good odour); P-O- (bad port and bad odour).

**Table 1.** Best sets of encoding parameters for describing remote episodic memory precision. Best doublets and triplets of encoding variables selected by using a descriptive linear regression model to explain remote episodic memory performance during the recall test (n = 24). The Bayesian information criterion (BIC) value was calculated for each regression. The lower the BIC value is, the better the linear regression model approximates memory performance. Only one parameter was recurrent in all selected models, the *Index CR_P+O-_E1.1,* which corresponds to the index of avoiding the unpleasant drinking solution in the P+O- configuration during the first E1 presentation. *CR* (correct rejection; avoiding drinking from the incorrect configuration); nb (number); *FA* (false alarm; drinking from the incorrect configuration). E1.1 (first episode E1); E2.1 (first episode E2); E1.2 (second episode E1); E2.2 (second episode E2); P+O+ (good port and good odour); P+O- (good port and bad odour); P-O+ (bad port and good odour); P-O- (bad port and bad odour).

|  | 1st variable | 2nd variable | BIC value |
| --- | --- | --- | --- |
| 1<sup>st</sup> doublet | **Index CR_P+O-_E1.1** | Index CR_P+O-_E2.2 | -22.07 |
| 2<sup>nd</sup> doublet | **Index CR_P+O-_E1.1** | Visits nb_P+O-_E2.2 | -22.01 |

***Table 1. Best sets of encoding parameters for describing remote episodic memory precision***
|  | 1st variable | 2nd variable | 3rd variable | BIC value |
| --- | --- | --- | --- | --- |
| 1<sup>st</sup> triplet | **Index CR_P+O-_E1.1** | Visits nb_P+O-_E2.2 | FA nb_P+O-_E1.2 | -24.23 |
| 2<sup>nd</sup> triplet | **Index CR_P+O-_E1.1** | Licks nb_P+O-_E1.2 | Visits nb_P+O-_E2.2 | -22.92 |
| 3<sup>rd</sup> triplet | **Index CR_P+O-_E1.1** | Visits nb_P-O+_E1 | Licks nb_P-_E1.2 | -22.68 |

To emphasize the importance of behaviour during this critical initial episodic experience, we selected a set of behavioural parameters for each encoding session that best predicted remote memory performance (Fig 5b). Regardless of the number of parameters used, the best model (lower BIC value) was obtained with behavioural parameters from the first Episode 1. Finally, we explored the relative importance of parameters related to pleasant (sugar) and unpleasant (quinine) experiences by investigating (within each episode) the behavioural parameters associated with each type of drinking solution in the good port P+ (Fig. 5c). The model scores showed that across encoding episodes, unpleasant experiences were better predictors of remote episodic memory precision than pleasant experiences. Importantly, the beneficial experience of the first episode was not related to the odour exploration level since the odour sampling times of different memory profiles were similar during this encoding session (Fig 5d) (*P*_Kruskall-wallis_>0.05).

These results show for the first time that individual unpleasant experiences during nontraumatic life episodes strongly influence the content and precision of remote episodic memories.

### Brain networks associated with remote episodic memories differ according to memory content and level of detail

Brain networks associated with the consolidation and retrieval of remote episodic memories are critical in the memory field. Few studies have explored this issue in either humans or animals, and only a few tasks produce remote episodic memories by using encoding conditions similar to everyday life. Here, we used cellular imaging of immediate early genes (IEGs) after retrieval in various brain areas to determine what regions are recruited during recall of remote episodic memory as a function of their content and level of detail (Supplementary Fig. 3). We analysed the expression of *c-Fos*, which maps neural ensembles activated by experience, and *Zif268*, which is required for synaptic plasticity and long-term memory^32–34^. IEG expression was investigated in rats that exhibited *WWW* or *Where* profiles during recall (Fig. 3) and compared to control rats that experienced only routine sessions^30^.

For *Where* rats that remembered place information, we observed significantly increased c-Fos and Zif268 expression in 6 brain areas, including the lateral and dorso-lateral parts of the orbitofrontal cortex (LO.DLO) (Zif268 *P*=0.025), anterior part of the retrosplenial cortex (aRSG) (c-Fos *P*=0.025), habenula (Hb) (Zif268 *P*=0.0039) and dorsal hippocampus (c-Fos: dDG *P*=0.016; Zif268: dCA1 *P*=0.037, dCA3 *P*=0.025) (Fig. 6a, c). Olfactory areas, the prefrontal cortex, the basolateral amygdala, and the parahippocampal and ventral hippocampal regions were not recruited in *Where* rats during the remote recall test.

**Figure 6.**
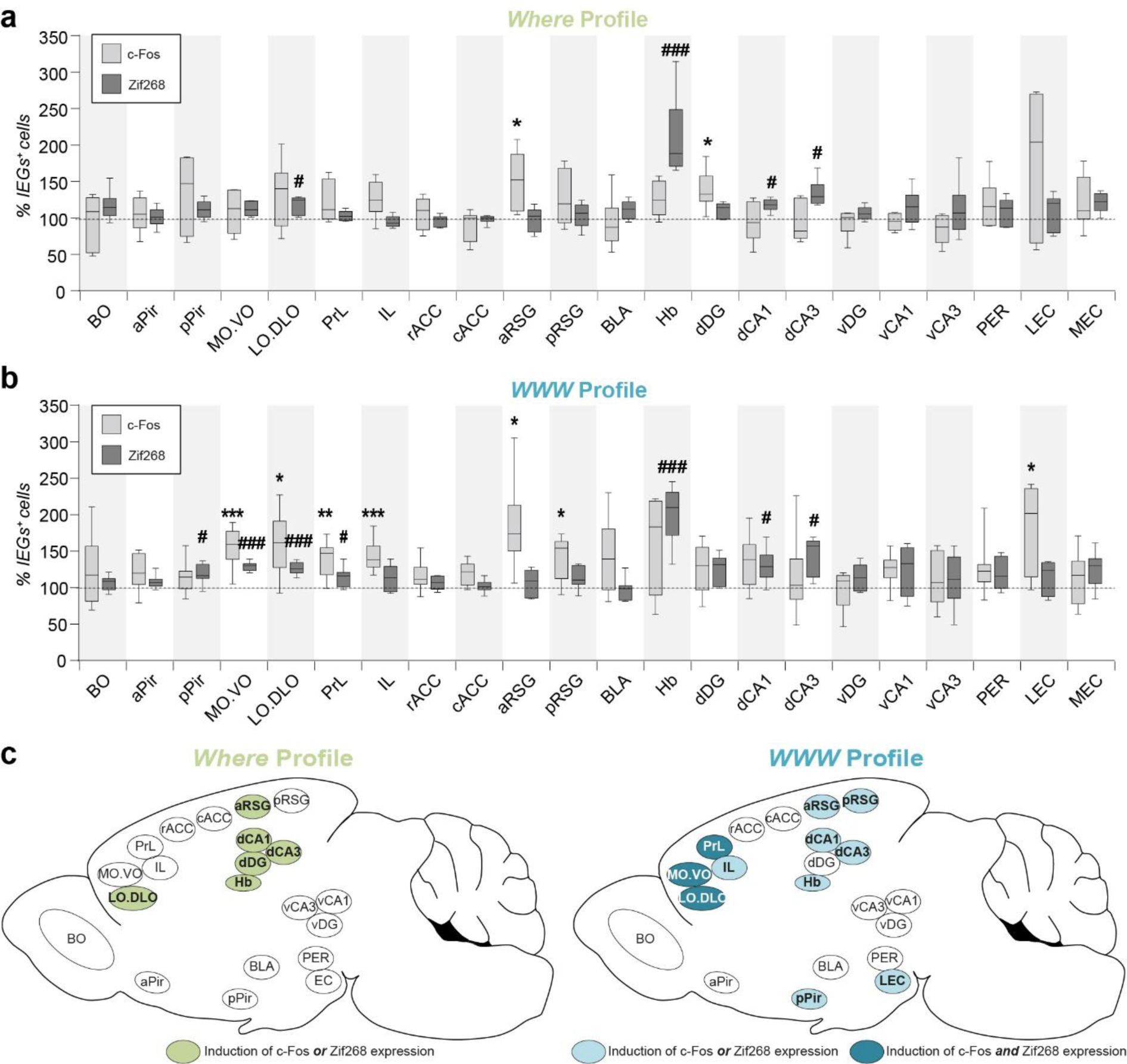
Different brain networks recruited for remote episodic memory with various content. Density of c-Fos^+^ (light grey) and Zif268^+^ (dark grey) cells in analysed brain areas for **(a)** *Where* and **(b)** *WWW* profiles after the recall test. Data were normalized and statistically compared to the control group (value of 100%). **(c)** Summary diagram representing brain areas recruited for *Where* and *WWW* rats. In *Where* rats that remembered only location information in the context, c-Fos or Zif268 expression was significantly increased in the LO.DLO, aRSG, Hb, dCA1, dCA3, and dDG regions. In contrast, *WWW* rats that recollected the entire episode recruited larger brain networks. Some of these rats recruited brain areas that were similar to those recruited by *Where* rats, including aRSG, Hb, dCA1, and dCA3. New areas were also recruited, including IL, pRSG, pPir, and LEC, and c-Fos and Zif268 expression was induced in several cortical regions, including MO.VO, LO.DLO and the PrL cortex (dark blue). Mann‒Whitney U tests for statistical comparisons between groups (n*_Control_* = 6, n*_WWW_* = 9, n*_Where_* = 6). *p<0.05, **p<0.01, ***p<0.005 for c-Fos; #p<0.05, ##p<0.01, ###p<0.005 for Zif268. OB (olfactory bulb); aPir/pPir (anterior and posterior piriform cortices); MO.VO (medio-ventral), LO.DLO (dorso-lateral) part of the orbitofrontal cortex; PrL (prelimbic) and IL (infralimbic) cortices; rACC, cACC (rostral and caudal anterior cingulate cortices); aRSG, pRSG (anterior and posterior retrosplenial cortices); BLA (basolateral amygdala); Hb (habenula); dDG, vDG (dorsal and ventral dentate gyrus); dCA1, dCA3 (dorsal) and vCA1, vCA3 (ventral) hippocampus; PER (perirhinal cortex); LEC, MEC (lateral and medial entorhinal cortices).

In contrast, in *WWW* rats that recollected the entire episode, different and larger brain networks were recruited, including a hard core of cortical regions in which both IEGs were induced (Fig. 6b, c), as in all parts of the orbitofrontal cortex (c-Fos: MO.VO *P*=0.0047, LO.DLO *P*=0.034; Zif268: MO.VO *P*=0.0014, LO.DLO *P*=0.0022) and the prelimbic cortex (PrL) (c-Fos: *P*=0.0067; Zif268: *P*=0.025). More brain regions with significantly increased c-Fos or Zif268 expression were observed in *WWW* rats than in *Where* rats, including the posterior piriform cortex (pPir) (Zif268 *P*=0.018), infralimbic cortex (IL) (c-Fos *P*=0.0032), posterior retrosplenial cortex (pRSG) (c-Fos *P*=0.045) and lateral entorhinal (LEC) cortex (c-Fos *P*=0.014). Furthermore, certain brain regions were activated for both memory profiles, including the anterior retrosplenial cortex, habenula and dorsal hippocampus (c-Fos: aRSG *P*=0.014; Zif268: Hb *P*=0.0014; dCA1 *P*=0.013, dCA3 *P*=0.013). Interestingly, different brain networks were activated in *WWW* rats during memory recall after one^30^ and 30 days, suggesting that episodic memory engrams transform over time (Supplementary Fig 4). While recall of a recent episodic memory recruits the entire hippocampus, only the dorsal part remains engaged for remote memories, with IEG expression restricted to Zif268. Moreover, cortical memory traces are reorganized, with the OFC critical for recalling remote memory and the rACC critical for recalling recent memory^30^. Finally, the posterior piriform and lateral entorhinal cortices, which are implicated in odour processing, are recruited only after 30 days of retention, suggesting the key role of odours in remote episodic memories.

Our findings reveal that brain networks associated with recall of remote episodic memory are intrinsically related to the content and level of detail in the memories. Incomplete episodic memories that include only context and spatial information recruit the CA1, CA3 and DG of the dorsal hippocampus and associated brain areas, such as the RSG. In contrast, complete remote episodic memory is associated with larger networks, including brain areas for olfactory processing (piriform and lateral entorhinal cortices), large cortical regions (orbitofrontal, prefrontal, and retrosplenial), the habenula and the CA1/3 regions in the dorsal hippocampus.

### Functional networks during recall can reflect the state and fate of a remote episodic memory

Memory recall mobilizes functional networks, with several interacting brain areas jointly activated^35, 36^. To assess coordinated activity among multiple brain regions, we calculated all interregional correlations among IEG^+^ cell densities across individual rats with *Where* and *WWW* profiles. We used significant Spearman coefficients (positive and negative) for all pairs of structures to construct correlation matrices and graph theory approaches to extract the functional organization of each memory network.

c-Fos expression graphs of the *Where* and *WWW* groups showed 30 and 23 significant correlations, respectively, with different coactivation patterns (Fig. 7a-b, left). The functional networks of *Where* rats were dense and homogeneously distributed within the brain (Fig. 7a, right) did not differ from a randomized functional network (Supplementary Table 1, global efficiency), suggesting a defect in coordinated brain activities in these rats during recall. Moreover, *Where* (27%) rats had twice as many negative correlations (dotted lines) as *WWW* rats (13%). Negative correlations in *Where* rats were all highly significant in terms of strength and concentrated in two areas, namely, dDG-vCA1-aPIR-pPIR-LEC and vCA3-PrL-IL. Interestingly, dDG activity was negatively correlated with many olfactory areas, such as aPir, pPir and LEC, which could explain why *Where* rats no longer associated odour information with other memory components. In *WWW* rats (Fig. 7b, right), while most correlations were observed in olfactory (30.4%) and cortical (41.3%) areas, coactivations were nearly absent in the hippocampus (4.3%), except between dCA1 and MO.VO. We applied three centrality measures and bootstrap statistical analyses to identify the central nodes in each network (Supplementary Table 2a). While dCA3 and MEC were the only major regions in the networks of *Where* rats, recall in *WWW* rats was supported by several olfactory processing regions, including the olfactory bulb (CG, PGL), aPir, MEC and IL and pRSG cortical areas.

**Figure 7.**
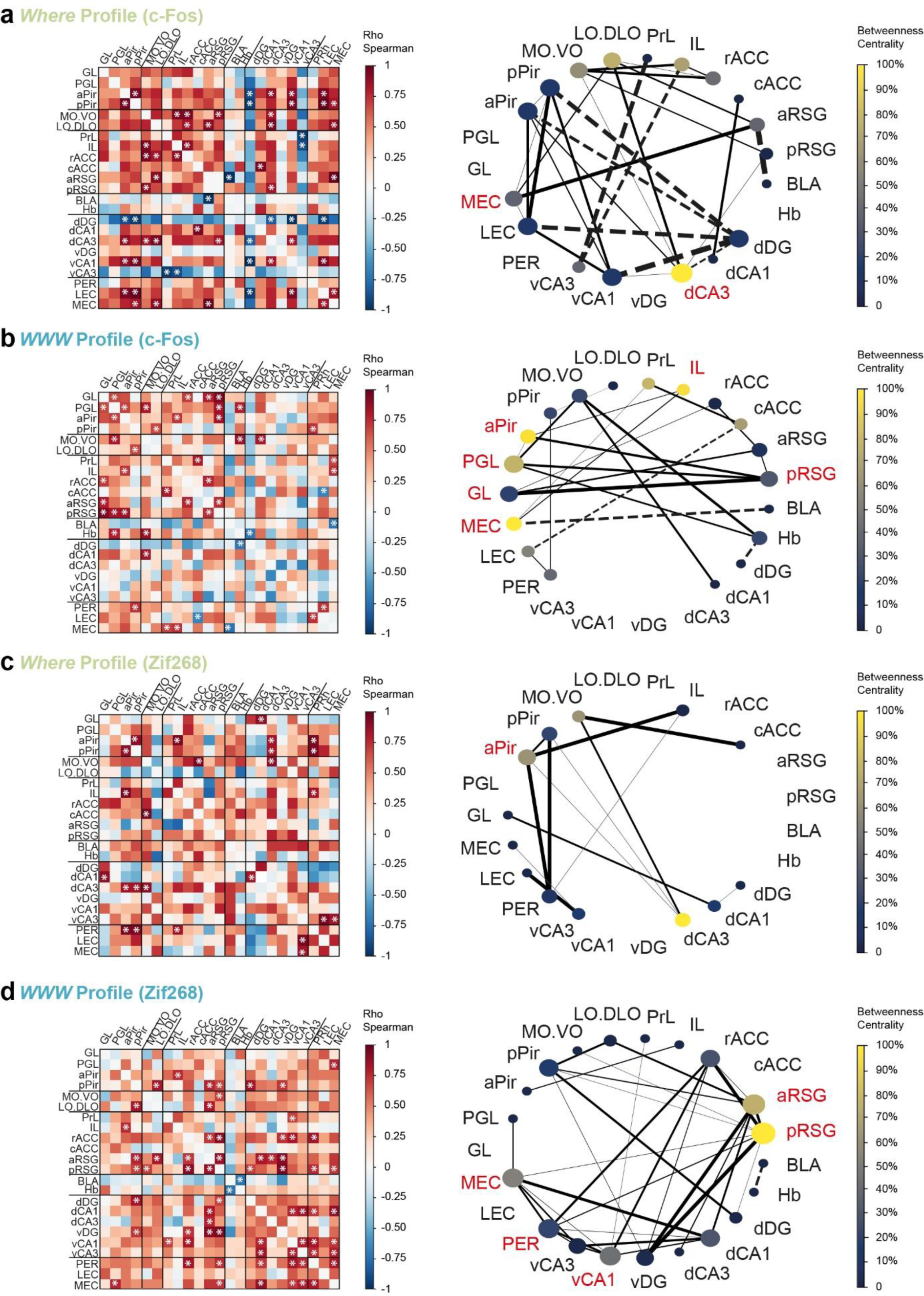
The state and fate of remote episodic memory according to functional network analyses. **(a-d, left)** Interregional correlation matrices for each memory profile (*Where* and *WWW* rats) according to c-Fos (a, b) or Zif268 (c, d) expression. The X and Y axes correspond to different brain areas. The colour code reflects the correlation strength (scale of Rho Spearman coefficient). Significant correlations (p_Spearman_≤0.05) are indicated with a star (n*_WWW_* = 9, n*_Where_* = 6). **(a-d, right)** Graphs were generated by considering only significant correlations, including both positive (full lines) and negative (dotted lines) correlations. The thickness of the connections is proportional to the correlation *strength*, the size of the node is related to the *degree*, and the node colour reflects the *betweenness centrality* (colour scale). Network hubs are highlighted in red. **(a, b)** The c-Fos graphs were different in *Where* and *WWW* rats. The *Where* network was dense and homogeneously distributed in the brain, and there were more negative correlations and few hubs (only dCA3 and MEC). In *WWW* rats with complete remote episodic memory, significant correlations were observed in olfactory and cortical areas, whereas coactivation was restricted to the dCA1 of the hippocampus with MO.VO. Five hubs were obtained in the *WWW* graph: GL and PGL of the OB, pPir, pRSG, IL and MEC. **(c, d)** The Zif268 graphs were considerably different between *Where* and *WWW* rats, with the opposite pattern. In *Where* rats, most coactivations were observed in the MO.VO, olfactory and parahippocampal areas, and the aPir was the only hub. In *WWW* rats, functional connectivity was mainly observed between cortical and medial temporal lobe areas, and five hubs were observed: the entire RSG, vCA1, MEC and PER. GL (granular layer of the olfactory bulb); PGL (periglomerular layer of the olfactory bulb); aPir/pPir (anterior and posterior piriform cortices); MO.VO (medio-ventral), LO.DLO (dorso-lateral) part of the orbitofrontal cortex; PrL (prelimbic) and IL (infralimbic) cortices; rACC, cACC (rostral and caudal anterior cingulate cortices); aRSG, pRSG (anterior and posterior retrosplenial cortices); BLA (basolateral amygdala); Hb (habenula); dDG, vDG (dorsal and ventral dentate gyrus); dCA1, dCA3 (dorsal) and vCA1, vCA3 (ventral) hippocampus; PER (perirhinal cortex); LEC, MEC (lateral and medial entorhinal cortices).

The same analyses were performed for Zif268. In in contrast to the c-Fos connectome, there were no negative correlations and 2.6 times more significant coactivations (34 *versus* 13) in *WWW* rats than in *Where* rats (Fig. 7c-d, left). Moreover, functional networks (Fig.7c-d, right) and hubs (Supplementary Table 2b) are diametrically opposed between groups. In *Where* rats, most coactivations involved MO.VO, olfactory areas, and parahippocampal regions, and only the aPir was a hub (Fig. 7c, right). In contrast, in *WWW* rats, most coordinated activities were observed between cortical and medial temporal lobe areas, and five hubs were highlighted, including the entire retrosplenial cortex, vCA1 and some parahippocampal areas (MEC, PER) (Fig. 7d, right).

We used two IEGs to map brain networks activated during recall because their expresison may have distinct functional implications. The c-Fos connectome was used to assess network activity that reflected the state of memory during recall, with less specific and efficient brain networks developed for incomplete memories (*Where* rats) than for complete remote memories (*WWW* rats), for which olfactory and cortical areas were preferentially recruited. In contrast, given its critical role in synaptic plasticity and memory consolidation and reconsolidation^32–34^, the Zif268 connectome could reflect the fate of post-recall memories. While *Zif268*-related synaptic plasticity processes were observed in olfactory and parahippocampal areas in *Where* rats that did not associate odours with episodes, *Zif-268*-related synaptic plasticity reinforcement was induced between cortical and hippocampal areas in *WWW* rats that remembered the entire episode. This result suggests that during remote memory recall, parts of the networks are activated in response to remote episodic memory expression, whereas in other parts, synaptic plasticity processes are induced and may be related to strengthening and updating memory traces.

### The emotional brain has a central role in maintaining accurate remote episodic memories

Brain networks recruited during recall and their evolution are related to the detail levels of remote episodic memories. However, because different remote memories of the same life episode contain overlapping information, it can be difficult to determine how a complete memory is recollected. Therefore, we examined which brain areas are specifically activated by retrieval of an entire episode by directly comparing the IEG expression of *Where* and *WWW* rats.

Only 6 brain areas showed more c-Fos^+^ or Zif268^+^ cells in *WWW* rats than in *Where* rats (Fig. 8 a-c and Supplementary Fig. 5): the medial part of the OFC (both IEGs; c-Fos: MO.VO *P*=0.013; Zif268: MO.VO *P*=0.007), infralimbic cortex (Zif268=0. 025), caudal ACC (c-Fos *P*=0.013), BLA (c-Fos *P*=0.045) and entire hippocampal CA1 area (c-Fos dCA1 *P*=0.045, vCA1 *P*=0.0095) (Fig. 8e). We next analysed whether the activation level of these brain areas was correlated with memory precision during recall. These correlations were observed in only 3 brain regions involved in the valence and emotional processing of information: the MO.VO (c-Fos: Rho=0.52, *P*=0.05; Zif268: Rho=0.8, *P*=0.003), BLA (c-Fos Rho=0.72, *P*=0.008) and vCA1 (c-Fos Rho=0.63, *P*=0.02) (Fig. 8d). These correlations were not related to the level of reinforcement received during the test since no links between activated brain areas and the total number of licks, which reflected the amount of water consumed, were observed (Supplementary Fig. 6). This result reinforces the findings of our computational study, showing the critical influence of individual emotional experiences during episodic encoding on remote memory precision. In addition, these regions were related to each other since they showed positively correlated coactivation, except for MO.VO with BLA (Supplementary Fig. 7). This result supports the idea of a core network of interconnected brain areas that all contribute to accurate recollection of remote episodic memory.

**Figure 8.**
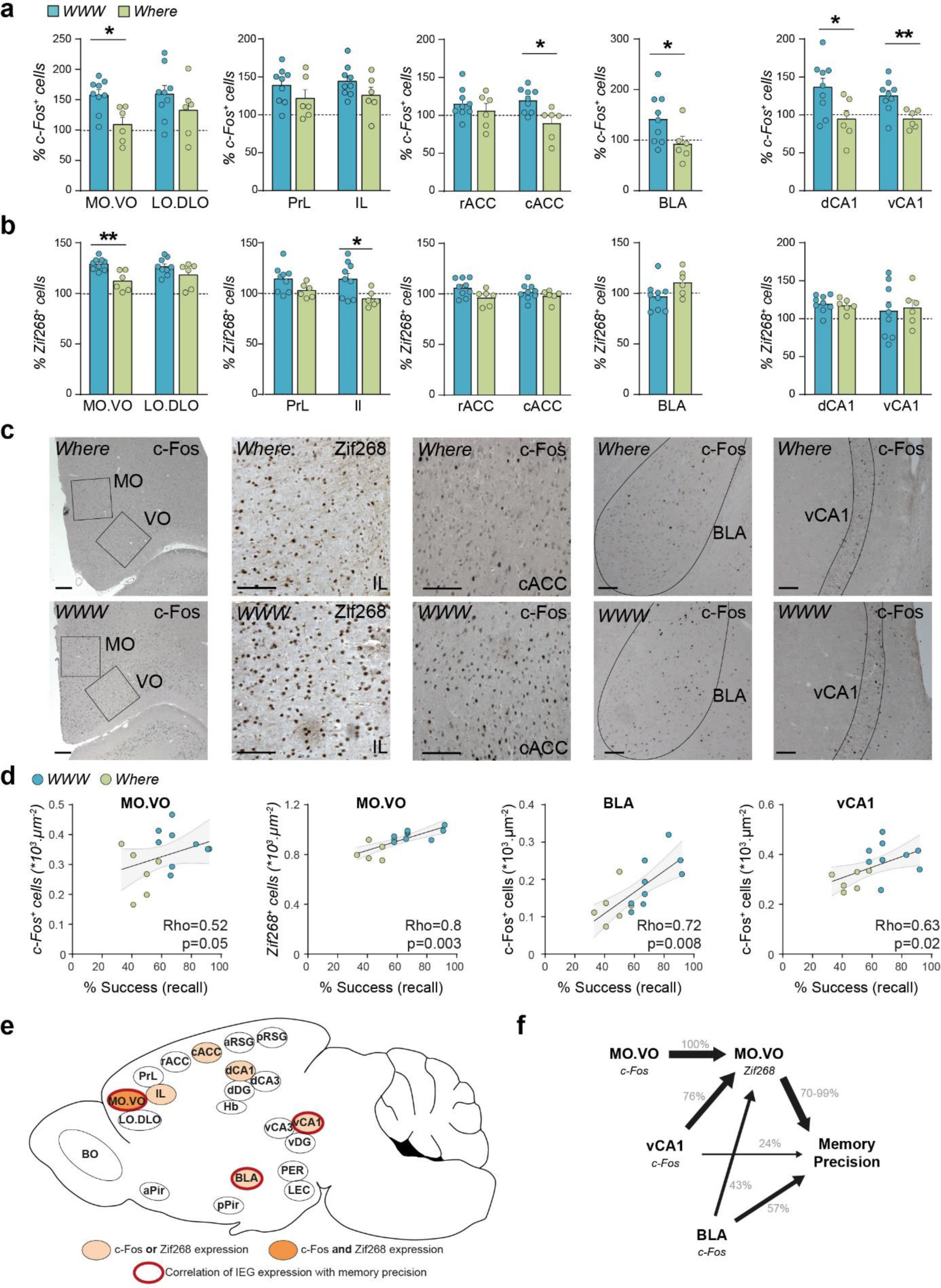
The brain emotional network contributing to the maintenance of an accurate remote episodic memory. **(a, b)** Statistical comparisons of the density of c-Fos^+^ and Zif268^+^ cells in *WWW* and *Where* rats, reflecting brain areas specifically recruited for accurate remote episodic memory. Data were normalized to the control group (value of 100%). The expression of one of the two IEGs was induced in five brain areas in *WWW* rats, namely, the cACC, BLA, dCA1, vCA1 and IL, whereas the cell density of both c-Fos and Zif268 was increased in the MO.VO of the OFC. **(c)** Photomicrographs showing the induction of IEG expression in *WWW* and *Where* rats. Scale bar 100 µm. **(d)** Positive Spearman correlations between IEG^+^ expression and remote episodic memory precision (success rate during recall) in the MO.VO, BLA and vCA1 of *WWW* and *Where* rats. **(e)** Summary diagram representing brain areas specifically recruited in *WWW* compared to *Where* rats (light and dark orange background) and regions in which IEG expression was directly correlated with memory precision (thick red outline). **(f)** Schematic representation of the causal mediation analysis performed on the emotional core network that contributed to accurate remote episodic memories. Zif268 expression in the MO.VO was the main direct mediator of remote memory precision and the influence of c-Fos-induced expression in the MO.VO, vCA1 and BLA. BLA recruitment was a second direct mediator of accurate episodic recollection, whereas the direct influence of the vCA1 was not significant. Group data are represented as the mean ± SEM (n*_Control_* = 6, n*_WWW_* = 9, n*_Where_* = 6). * p_Mann- Whitney_< 0.05; ** p_Mann-Whitney_<0.01, Mann‒Whitney U tests for statistical comparisons between the *Where* (n = 6) and *WWW* (n = 9) profiles. OB (olfactory bulb); aPir/pPir (anterior and posterior piriform cortices); MO.VO (medioventral), LO.DLO (dorso-lateral) part of the orbitofrontal cortex; PrL (prelimbic) and IL (infralimbic) cortices; rACC, cACC (rostral and caudal anterior cingulate cortices); aRSG, pRSG (anterior and posterior retrosplenial cortices); BLA (basolateral amygdala); Hb (habenula); dDG, vDG (dorsal and ventral dentate gyri); dCA1, dCA3 (dorsal) and vCA1, vCA3 (ventral) hippocampus; PER (perirhinal cortex); LEC, MEC (lateral and medial entorhinal cortices).

Finally, we applied causal mediation analyses to determine the mechanisms underlying the relationships between these entities (Fig. 8f and Supplementary Table 3). The principal mediator of memory precision was Zif268 expression in the MO.VO (70-99% direct effect), which is also influenced by the activation of the MO.VO (c-Fos MO.VO 100% *P*=0.01), vCA1 (76% *P*=0.02) and BLA (43% *P*=0.02). BLA activation appears to be a second potential mediator of remote memory precision (57% *P*=0.09 for direct effect and *P*=0.08 for mediated effect), whereas the direct influence of vCA1 was not significant (24% *P*=0.58).

Our results show that the network activated by recollecting old nontraumatic episodic memories includes three brain areas (OFC-vCA1-BLA) involved in processing emotions and information value. Synaptic plasticity processes in the medio-ventral OFC might play a primary role in maintaining the engrams of precise remote episodic memories.

## Discussion

Our first aim was to explore the characteristics of remote episodic memory in rats by using a task that models the attributes of episodic memories in humans. Although several studies have explored episodic memory in animals^3, 14–24^, none have provided data on old, complete episodic memory formed in an incidental and occasional manner. Our task is more controlled than autobiographic memory studies, more natural and immersive than laboratory approaches used in human studies, and forms memories more incidentally than most animal paradigms, which use extensive training. Moreover, our task allows remote episodic memory and their intrinsic variability to be accessed directly. Direct access to remote episodic memories that are *i)* controlled in terms of information content, *ii)* less semantic by reducing the number of repetitions during encoding and *iii)* ingrained within initial immersive contexts is a major challenge in human and animal studies that aim to explore the engrams of episodic memories that remain clear over time^5, 6, 37–40^.

In this study, we show that a majority of rats formed a robust remote episodic memory after experiencing only two life episodes with different combinations of contextual, spatial and olfactory information. Similar to humans^41, 42^, we found that rats naturally displayed not one but several remote long-term episodic memories with distinct content and detail levels. The individual profiles are highly reproducible, and their variability increases over time, demonstrating that although they are robust, remote episodic memories are also malleable and sensitive to retrieval interference, which are common in daily life when examining information in different episodes. Because they are the first information to be encoded and the most explored information in our task, the contextual and spatial information are preserved best over time. Odours are the last information to be encoded and the least explored during episodes (4 min maximum in each 40 min episode). However, odours are critical components in emotional, detailed and old memories in humans^42–44^, and our results show that rats have an entire remote episodic memory if they experienced unpleasant experiences related to odours during the very first episode. In addition to reinforcing the idea that odours are critical components of episodic memories, this result is the first to provide evidence of the direct influence of individual experiences during episode encoding and the emotional value of information on the retention and accuracy of nontraumatic episodic memories. The origin of individual differences in recall during daily life should be determined in animals and humans^45^ because this is a major characteristic of this very personal memory and could thus be a step forward in understanding its subtleties.

The present study is the first to provide data on the supportive networks of remote episodic memory in animals. We show that the networks associated with episodic recall are intrinsically related to the content of remote episodic memories since recruited brain areas are representative of the stored information. The brain networks recruited in *Where* rats, that show an incomplete memory, are more restricted and less efficient in terms of functional connectivity and specific to networks of memories with contextual and spatial information (dorsal hippocampus, retrosplenial cortex, and habenula)^46–50^. In contrast, *WWW* rats that remember all episodic information recruit larger brain networks, including the same spatial network and brain regions that are functionally connected and implicated in olfactory memory (piriform, lateral entorhinal, orbitofrontal, prelimbic and infralimbic cortices)^51–56^. A comparison of the *Where* and *WWW* profiles reveals that complete recollection of remote episodic memory recruits a core network of interconnected brain regions (medio-lateral orbitofrontal, ventral CA1 and basolateral amygdala)^57–62^ that are implicated in the valence and emotional processing of information and directly correlated with memory accuracy during recall. This novel result provides new perspectives for exploring the critical role of the emotional brain in old nontraumatic episodic memories and the specific role of odours and their value in maintaining memory of personal events^42–44^.

Our data contribute to the still debated topics of the neuronal substrate of episodic memory and its fate over time, and our results reinforce the idea that the nature and quality of a memory matter more than its age^63, 64^. In recent years, various models have been proposed reconcile data collected on this issue. Our results support the posterior medial–anterior temporal (PMAT)^65–67^ model, with the AT system gathering regions that are functionally connected to the anterior hippocampus (amygdala, orbitofrontal and perirhinal cortices) and process perceptual, semantic, and emotional memory components as general information and the PM system regrouping brain regions that are functionally connected to the posterior hippocampus (as parahippocampal, retrosplenial, posterior cingulate and medial prefrontal cortices) and process contextual, fine-gained and detailed information. Our data also support multi-trace theory (MTT) and trace transformation theory (TTT) models, which postulate that in contrast to semantic memories, episodic memories depend on the reactivation of memory traces within the hippocampus, while their transformation over time and various experiences lack details and context specificity and are thus better represented in distributed neocortical networks^10, 12, 40, 63, 64^. However, detailed episodic or context-specific memories could depend on the hippocampus. Consistent with these models, we have previously shown that recent episodic memory is supported by the entire hippocampus and a large cortical motif^30^. Here, we show that for remote episodic memories, only the dorsal hippocampus remains engaged as the involvement of the orbitofrontal cortex increase (Supplementary Fig. 4). In addition, our results show that the orbitofrontal cortex hosts synaptic plasticity mechanisms that correlate with the precision of remote memory; thus, this region is critical for the retrieval of old and complete episodic memory. This finding is consistent with the major role of the orbitofrontal cortex in the processing and relevance of associative olfactory memory^27, 57–61^ and the idea that odours and their emotional value are crucial for remote episodic memory^43, 44^. The involvement of the hippocampus in remote episodic memories in both *Where* and *WWW* rats indicate that their respective memories are still associated with their context, regardless of their content. However, because only *Zif268,* an IEG mainly implicated in synaptic plasticity^32–34^, is expressed in the dorsal hippocampus during recall, our results support the recent scene (re)construction theory, with neocortically consolidated elements reconstructed into new hippocampal traces of sequences of scenes that comprise the past event^39^. Finally, our results support the idea that remote episodic memories remain highly dynamic and malleable since, during retrieval, engrams are either updated for rats that no longer associate all information (*Where* rats) or reconsolidated in rats that still remember the entire episode (*WWW* rats). This finding suggests that even if memory content changes over time, reactivation could be sufficient to strengthen, restore or update remote episodic memories.

Overall, our data provide new perspectives on the origin of individual differences in remote episodic memory and the impact of emotions, odours, and reactivations on its fate and evolution over time. Our animal model offers the outstanding opportunity to elucidate mechanistic aspects related to brain networks and plasticity that support the maintenance and transformation of episodic memories over time. The ultimate challenges are to determine the functioning of episodic memory which is so important for individuals, and to better understand its alterations in certain disorders to address them effectively^68^.

## Methods

### Animals

All experiments were performed in accordance with the European Directive (2010/63/EU) and the French Ministry of Higher Education, Research and Innovation (n°APAFIS #7201- 2016101213348550). A total of 40 male Long Evans rats (Charles River Laboratories, Italy) that were 8 weeks old (296 ± 22 g) at the beginning of the experiment were housed in groups of 2 in standard cages under a 12 h light/dark cycle. The experiments were performed during the light phase. Food was available *ad libitum*, but water consumption was controlled by gradually reducing daily access to two 40 min sessions (8 am and 6 pm). The weight of the animals was monitored weekly, and the amount of water consumed was monitored daily to ensure the welfare of the rats throughout the experiment.

### The EpisodiCage

The experimental arena and details of the behavioural protocol have been extensively described in previous papers by our group^30, 31^. Briefly, the experimental arena is a rectangular area equipped with 4 odour ports associated with a retractable drink-dispensing pipette (Fig. 1a, left panel). Two olfactometers allow the occasional diffusion of odours, which remain confined within the port, through a continuous vacuum system. When a rat visits a port, if a nose poke is detected, an odour is immediately diffused for 10 sec; 3 sec after the nose poke, the pipette delivers a drinking solution for 10 sec; then, the pipette retracts (Fig. 1a, right panel). Rats explored the arena and ports freely and at their own pace. Visits to the ports and licks made by the rats on the pipettes were recorded and temporally reconstructed with homemade *Neurolabscope* software, and the global behaviour was recorded with five cameras and analysed by the homemade video tracking software *Volcan*. The floor of the arena was removable, and its appearance and texture can be changed. Loudspeakers and a video projector were used to provide a multisensory environment for episodes with different contexts.

### Odorants and drinking solutions

The odorants (Sigma‒Aldrich, France) used were geraniol (#163333) and eugenol (#E51791) in the shaping phase and D-carvone (#435759)/isoamyl acetate (#W205508) and trans-anethole (#117870)/citral (#W230308) in the episode sessions. Depending on the task phase, the pipettes delivered water, sugar solution (sucrose 6%, Sigma‒Aldrich, France) or a bitter solution (quinine 0.06%, Sigma‒Aldrich, France).

### Episodic memory task

#### Shaping

During 10 daily sessions (15-20 min or a maximum of 20 port visits) in a neutral environment in an EpisodiCage with all active ports (Fig. 1b-c), rats learned that a visit with a nose poke in a port induced a pipette that delivered water for a limited time. During the first 6 daily sessions, no odours were diffused. During the next 4 daily sessions, rats were habituated and encountered two different odour stimulations at the ports that were associated with either a sweet or bitter drinking solution. A delay of 3 sec between the nose poke and the pipette delivery was employed to ensure that the rat associated the odours with the drinking solution. The odours presented during the shaping phase were not used after this phase.

#### Routine sessions

Between the shaping phase and the episodes, the rats experienced routine sessions in a neutral environment with 4 active ports and no odour for 3 days (20 min or a maximum of 24 port visits) (Fig. 1c). This step enhances the salience of the following episodes and serves as a baseline for the various behavioural analyses.

#### Episode exposure

During a 40 min maximum session (or 24 port visits), rats experienced two distinct life episodes (Episode 1/E1 and Episode 2/E2) (Fig. 1b, d), allowing them to form “*What-Where-Which Context*” memory. To maximize the number of rats that remembered all episodic information without affecting the strength of the episodic memory itself^30^, each episode was repeated once, with 1 day between each exposure. Each episode was characterized by a salient event within the arena, which was transformed into a multisensory environment in which animals associated odour-port context information (Fig. 1d). During Episode 1, the environment was dark, with a rough grey floor and a video projection of a black and white checkerboard. Sounds of crickets at night were played. Only ports 2 and 3 (P2 and P3) were active, and two odours were associated in this episode (O1 and O2). During their free but limited exploration in time, rats experienced that O1 gives access to a pleasant sugar solution only on P2, whereas O2 on this port and both odours on P3 give access to an unpleasant bitter solution. During Episode 2, the environment was bright, Legos and pebbles were placed in the centre of the arena, the arena had a smooth black floor, and piano music was played. In this episode, only ports 1 and 4 (P1 and P4) were active, and two other odours were present (O3 and O4). Only O3 on P4 was associated with a sugar solution, whereas O4 on P4 and both odours on P1 led rats to experience an unpleasant bitter solution. During their spontaneous and undirected exploration, rats incidentally encoded that in each environment (*Which Context*), pleasant and unpleasant experiences were associated with specific odours (*What*) and specific port locations (*Where*). Each episode included four place-odour configurations: P+O+ (with sugar) and P+O-, P-O+, P-O- (with quinine).

#### Remote recall test

Remote episodic memory was tested 30 days after the episodes by placing the first group of animals (n = 19) in exactly the same situation as in Episode 2, except that only water was delivered to evaluate what the rats remembered (Fig. 1e). The recall test was restricted to 12 port visits or a maximum of 15 min to prevent extinction effects during the test due to the delivery of water and to compare behavioural and cell-imaging data based on the same experience level during recall^30^. A more challenging version of the test was also used for a second group of rats (n = 10) to examine whether recall of remote episodic memory was influenced by interference, namely, the simultaneous presentation of information from Episodes 1 and 2 (4-Port test, Fig. 4a). In this case, rats were placed in the Episode 2 context with the corresponding odours (O3 and O4) on ports P1 and P4 (*In context* configurations, *IC*), and the configurations of Episode 1 were also present (O1/O2 on P2/P3) (*Out of context* configurations, *OC*). This situation requires that rats sort their memories of E1 and E2 episodes to identify the elements (ports and odours) that were associated with the correct context^30^.

### Behavioural analysis

#### Visit and lick indices

The numbers of visits and licks performed by each rat for each configuration during each session were normalized by dividing the numbers of visits and licks by the total number of visits or licks during the session. These values were then averaged for each group of rats and are presented as the mean ± standard error of the mean (SEM).

#### Response types

For each visit during the test, the number of licks was expressed as binary data, namely, as “drink” or “avoid”, with a threshold of 10 licks serving as a minimum. This threshold was determined according to several experiments by extracting the average number of licks performed during each episode on configurations with quinine. Different types of common behavioural responses were then extracted according to traditional signal detection theory. When a rat drinks from the P+O+ configuration, it elicits a good response (HIT), whereas not drinking from this configuration is an omission error (MISS). For the other configurations (P+O-, P-O+, P-O-), a good response corresponds to a correct rejection (CR), whereas drinking corresponds to a false alarm (FA). These data were then normalized by calculating their proportion during each session and averaged for each group of rats (mean ± SEM). The total number of correct responses during each session was calculated by adding the HIT and CR scores, and the normalized success rate during each session was averaged across the experimental groups (mean ± SEM).

#### Individual performance and memory profiles

Given the heterogeneity of memory performance during recall observed for this task^30^, we extracted memory profiles by comparing the experimental data (visit, licks and response types) with theoretical data of “archetypal” rats. The classifications were performed blindly and compared among three experimenters. For the recall test performed in the same situation as Episode 2 (Figs. 1e and 3a), the different archetypes include: *What-Where-Which context* memory profile (*WWW*) for rats that drink mostly from the P+O+ configuration (licks index P+O+ = 1; P+O-/P-O+/P-O- = 0); *Where* memory profile for rats that remember only spatial information and drink mostly from the good port regardless of the two odours (licks index mainly on the good port: P+O+/P+O- = 0.5; P- O+/P-O-= 0); and *Indeterminate* memory profile for rats that drink indifferently from all configurations (licks index for each configuration: P+O+/P+O-/P-O+/P-O- = 0.25). During the 4-port recall test, the same profiles were determined according to the licks that were made in accordance with the E2 context (*In Context* configurations: P+O+, P+O-, P-O+, P-O-) and in concordance with the E1 context (*Out of Context* configurations: P+O+, P+O-, P-O+, P-O-) (Fig. 4 a, e).

#### Behavioural statistical analyses

The visit, lick and response type indices obtained for different configurations were statistically compared through nonparametric Friedman test and Wilcoxon rank test. Data from different groups of rats were compared through nonparametric Kruskal‒ Wallis test and Mann‒Whitney post hoc U test. The significance level was p≤0.05. All statistical analyses were performed using Systat® software.

### Multivariate regression analysis

Multivariate regression analyses were performed to determine whether individual behavioural variables observed during encoding are predictive of the precision of remote episodic memories. Twenty-four rats were investigated in this analysis. First, 420 encoding variables were selected, including the duration of odour exploration, the total number and indices of licks and visits, and the percentage of response types (HIT, MISS, CR, FA) for all possible configurations in each E1 and E2 session. Each encoding variable was individually correlated with the success rate during the recall test session. Redundant or irrelevant variables were excluded according to three criteria: *(i)* variables that did not vary within the experimental group and could not account for individual variability, *(ii)* variables that were absent in some rats, such as those corresponding to configurations not explored during the episodes, and *(iii)* variables that were strongly correlated with each other, which usually depend on each other and are thus redundant. In this case, when the Pearson correlation coefficient between two encoding measures was greater than 0.95, only the variable that was more correlated with the success rate during the recall test was retained. Finally, to prevent overfitting, a descriptive approach was used to select the best sets of variables explaining memory performance. The encoding variables were selected noniteratively in the linear regression model by systematically testing all possible pairs or triplets of encoding variables. The Bayesian information criterion (BIC) value was calculated for each possible regression. The lower the BIC value, the better the linear regression model approximates memory performance. Doublets or triplets of encoding variables that best explain memory performance were determined through this analysis. The multiple linear regressions and statistical analyses were performed with the Statsmodels module in Python 0.12.0 (https://www.statsmodels.org). The normality (Omnibus test and Jarque-Bera test), heteroscedasticity (Breush-Pagan test) and linearity (Harvey-Collier multiplier test) of the data were confirmed.

### Immediate early gene analysis

#### Experimental groups

The expression of *c-Fos* and *Zif268* IEGs in brain areas specifically recruited during remote episodic memory recall was analysed in rats that exhibited *WWW* (n = 9) or *Where* (n = 6) profiles during recall (Fig. 3) and compared to IEG expression in control rats (n = 6) that participated in only routine sessions (Fig. 1c), as previously described^30^. Control rats formed a remote memory of a familiar and stable environment with limited “*What-Where- In which context*” information since routine sessions were performed in a neutral context with no odours or salient information. Because of their behavioral heterogeneity and their insufficient number (n = 4), rats with an *Indeterminate* profile were excluded from the analysis (Fig. 3d).

#### Brain tissue preparation

Ninety minutes after the end of the recall test, rats were deeply anaesthetized with sodium pentobarbital (200 mg/kg) and transcardially perfused with a solution of 4% paraformaldehyde in 0.1 M phosphate buffer. The brains were removed and postfixed overnight at 4°C in the same solution, cryoprotected for 6 days in a solution of 30% sucrose in 0.1 M phosphate buffer and frozen (-30°C). Coronal serial sections (14 µm thick and spaced by 84 µm) from the olfactory bulb (Bregma 7.56 mm) to the posterior part of the hippocampus (Bregma -6.60 mm) were cut using a cryostat (LEICA CM1950, Leica Biosystems).

#### Immunohistochemistry of c-Fos and Zif268

The brain sections were preincubated in DAKO Target Retrieval Solution (Agilent) for 20 min at 95°C. After cooling for 20 min, the sections were treated with 0.5% Triton for 20 min, and endogenous peroxidases were blocked in 3% H_2_O_2_ (hydrogen peroxide 30%,). The sections were then incubated for 90 min in a blocking solution with 7.5% NGS (Normal Goat Serum, Jackson Immunoresearch), 2% BSA (Bovine Serum Albumin, Sigma‒Aldrich) and 0.1% Triton. The sections were incubated overnight at 25°C in the presence of a rabbit primary antibody specific for either c-Fos (1:3000, Santa Cruz

Biotechnology, #sc**-**52) or Zif268 (1:1000; Santa Cruz Biotechnology, #sc**-**189). After incubation for 2 hours at 25°C with biotinylated goat anti-rabbit secondary antibody (1:200; Eurobio), the sections were treated for 30 min with an avidin-biotin-peroxidase complex (1:200; Vectastain Elite ABC-HRP Kit, Eurobio). The revelation was performed in a solution of 3,3- diaminobenzidine-tetrahydrochloride (DAB, Sigma‒Aldrich), 0.03% NiCl_2_ and 0.06% H_2_O_2_. The DAB concentration was 0.066% for c-Fos (15 min) and 0.05% for Zif268 (4 min). Finally, the sections were dehydrated in graded ethanol and coverslipped in DEPEX (Sigma‒Aldrich). *<u>Quantification and mapping of c-Fos- and Zif268-positive cells</u>*. Quantitative analyses of c-Fos-and Zif268-positive cells were conducted using a stereology station (Mercator Pro, Exploranova) coupled to an optical microscope (Axio Scope.A1, ZEISS). The counts were determined by an experimenter blinded to the experimental conditions at 20x magnification in semiautomatic mode. Positive cells were identified in two sections spaced by 168 μm in 23 brain areas that were anatomically defined according to the Paxinos and Watson atlas (Paxinos and Watson, 1998) (Supplementary Fig 3). In the olfactory bulb (OB), counts were made on 3 slices spaced by 504 μm. The analysed brain areas included the granular (GC) and periglomerular layer (PGL) of the olfactory bulb (OB); anterior and posterior piriform cortices (aPir, pPir); orbitofrontal cortex (medial MO, ventral VO, lateral LO, dorso-lateral DLO); prelimbic (PrL) and infralimbic cortices (IL); rostral and caudal anterior cingulate cortices (rACC, cACC); anterior and posterior retrosplenial cortices (aRSG, pRSG); dorsal (dCA1, dCA3, dDG) and ventral (vCA1, vCA3, vDG) hippocampus; perirhinal cortex (PER); medial and lateral entorhinal cortices (MEC, LEC); and subcortical areas, including the basolateral amygdala (BLA) and habenula (Hb).

#### Data analysis and statistics

The density of c-Fos^+^ and Zif268^+^ nuclei was calculated per mm^2^, averaged in two sections and analysed for each brain area. The values were averaged for each experimental group (*WWW* versus *Where* memory profiles) and normalized to 100% of the control group ± SEM. Statistical comparisons between the different groups were performed with nonparametric Mann‒Whitney U test. Nonparametric Spearman correlations were used to identify correlations between the density of c-Fos^+^ or Zif268^+^ cells and memory performance during the recall test (% of success). Correlations between c-Fos and Zif268 expression in each brain area were also performed to examine brain coactivations of IEGs.

### Functional connectivity analysis

#### Correlation matrices

All interregional correlations of the IEG^+^ cell density were calculated for each memory profile (23 brain areas, 253 correlations in each experimental group). Colour-coded correlation matrices for the *WWW* and *Where* rats were generated by using Rho Spearman coefficients among all pairs of brain areas (Fig. 7, left).

#### Network graph construction and characterization

Brain networks were constructed by using positive and negative significant Spearman correlations (p≤0.05)^69, 70^ in each experimental group and characterized quantitatively with graph-theoretic measures (Fig 7, right). Each brain area was represented by a node with a given size, referred to as its *degree*, which reflects the number of brain regions coactivated with this node. All significant correlations of IEG densities between two brain areas are represented by connections between nodes, and the thickness of the connection is proportional to the strength of the correlation. Dotted connections represent negative correlations of activity between two nodes. One of the main purposes of graph theory is to identify central nodes, thus revealing their critical role in information flows within a network. To accomplish this, three measures of centrality were calculated for each node: the *degree*, the *strength* and the *betweenness centrality* (Supplementary Table 2). The *strength* of a node corresponds to the sum of all significant covariations in an area. The higher this value is, the more central an area in the network is. *Betweenness centrality* refers to the number of shortest paths between other nodes that the given node intersects. In the present study, this metric reflects the extent to which a brain region acts as an intermediary between other brain areas. While the *degree* and *strength* metrics reflect the static centrality of nodes, the *betweenness centrality* reflects the importance of nodes in communication within the network. Another purpose of graph theory is to characterize the efficiency of the entire network. Two measures were calculated to assess the efficiency: the *global clustering coefficient* and the *global efficiency* (Supplementary Table 1). The *global clustering coefficient* is measures network segregation based on the probability that two nodes are connected if they have a common neighbour. The *global efficiency coefficient* measures network integration and reflects the speed of information exchange within the network.

#### Software and statistical analysis

Correlation matrices and graph networks were generated with the graphpype package in Python (https://github.com/neuropycon/graphpype), Radatools software (http://deim.urv.cat/~sergio.gomez/radatools.php) and the open-source toolbox NeuroPycon (Meunier et al., 2020). All statistical analyses were performed by using bootstrap analyses (Supplementary Table 1) that randomly sample the data 100 times to compare each measure to those determined by random data.

### Mediation analysis

To better understand the impact of brain area activation on memory performance during the recall test, we performed a mediation analysis. This analysis allows us to determine the relative proportion of direct *versus* mediated effects of each brain area on memory performance. The mediation analysis was performed with the Mediation class from the Python statsmodels toolbox. For each brain area, we performed analyses between its activation and the memory performance using the activation of other areas as putative mediators (one mediation analysis per other area). The statistical significance of the results was estimated using a bootstrap method with 1000 repetitions (Supplementary Table 3).

The results show direct and indirect effects only when significant mediated effects are present (p < 0.05); otherwise, we noted only direct effects.

## Author contributions

A.A. performed the experiments, analysed the data and wrote the paper. N.F.T. designed the tools and analysed computational and mediation data. D.M. provided tools for functional network analyses. A.G. participated in the behavioural experiments and analyses. S.G. designed tools for acquiring and analysing behavioural data. B.M. designed the EpisodiCage and tools for acquiring behavioural data. M.T. designed the video tracking software for acquiring behavioural data. N.R. designed the behavioural experiments and analyses. A.V. designed the research, experiments, data analysis procedures and wrote the paper. All authors edited and approved the final version of the manuscript.

## Acknowledgements

This work was supported by CNRS, University Lyon 1 and a doctoral fellowship from “L’observatoire B2V des mémoires” to A.A. We are very grateful to S. Laroche and A.M. Mouly for scientific discussions about this study and their comments on this paper. We also thank P. Salin and M. Rosier for their help with functional network tools, S. Bouret for his advice on the mediation analyses and O. Jelassi for advice on animal care.

