## Supplementary figures for "Distinct brain networks for remote episodic memory depending on content and emotional value"

### Supplementary figure legends

**Supplementary FIGURE 1**

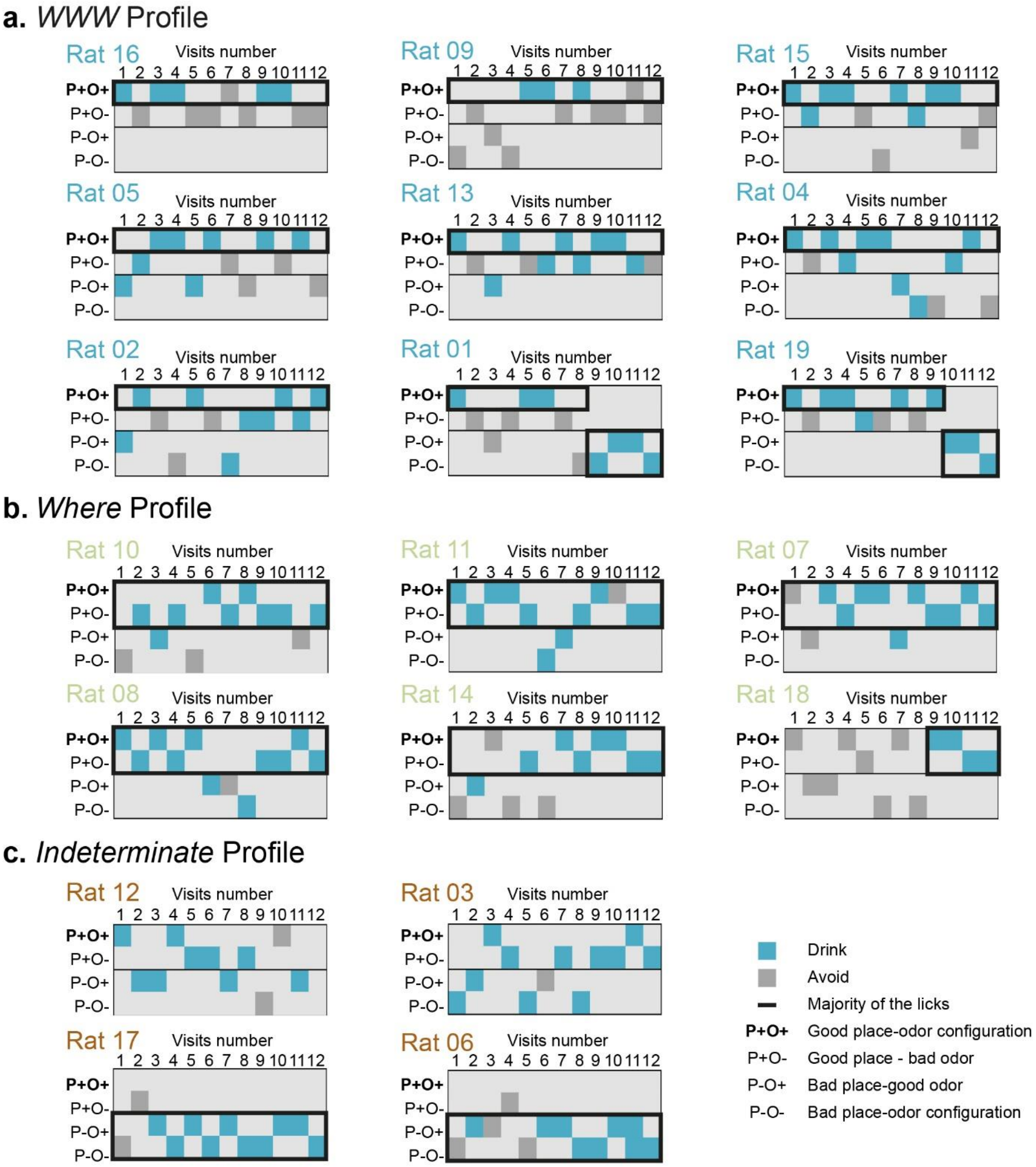


***Supplementary Figure 1. Robustness and stability of remote memory profiles during the recall test***

Individual and dynamic analyses during the recall test for each memory profile ($n_{WWW} = 9$, $n_{Where} = 6$, $n_{Ind} = 4$). For each rat, all visits (columns numbered from 1 to 12) are presented in front of the configuration encountered by the rat (lines P+O+, P+O-, P-O+, P-O-). A blue square indicates that a rat drank, and a grey square indicates that a rat did not drink from the given configuration. **(a)** *WWW* profile, with the black frame highlighting that rats predominantly drank from the P+O+ configuration and avoided other configurations during the recall test. **(b)** *Where* rats mainly visited and drank from the good ports (P+O+ and P+O-) during the session. **(c)** *Indeterminate* profile of two rats that drank from all configurations (Rats 12, 03) and two rats that mainly visited and drank from the wrong port (P-O+ and P-O-) during the recall test. P+O+ (good port and good odour); P+O- (good port and bad odour); P-O+ (bad port and good odour); P-O- (bad port and bad odour).

**Supplementary FIGURE 2**

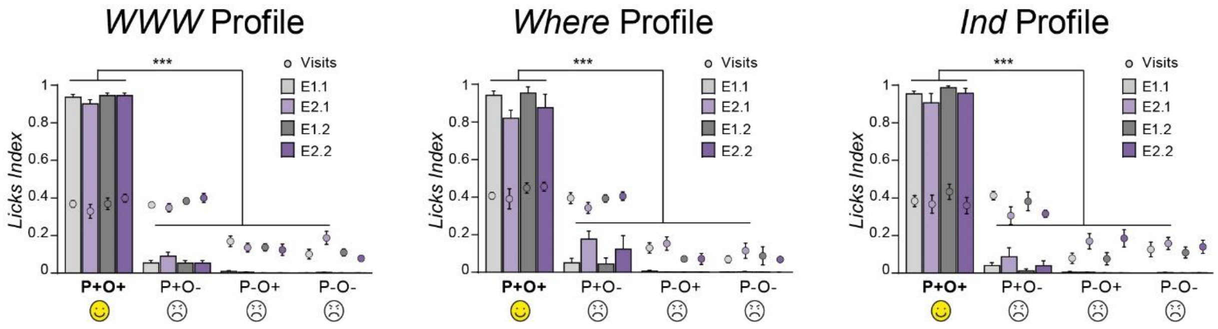


***Supplementary Figure 2. Remote memory profiles are not related to biased experiences during episode encoding***

Indices of licks and visits during episode encoding for each memory profile. Regardless of the profile, the rats visited and drank from the P+O+ configuration, which was associated with a pleasant experience of sugar solution, significantly more than the other configurations. Group data are expressed as the mean ± SEM ($n_{WWW}$ = 9; $n_{Where}$ = 6; $n_{Ind}$ = 4). ***p<0.005, Friedman tests followed by Wilcoxon tests for statistical comparisons between configurations. E1.1 (first episode E1); E2.1 (first episode E2); E1.2 (second episode E1); E2.2 (second episode E2). P+O+ (good port and good odour); P+O- (good port and bad odour); P-O+ (bad port and good odour); P-O- (bad port and bad odour).

**Supplementary FIGURE 3**

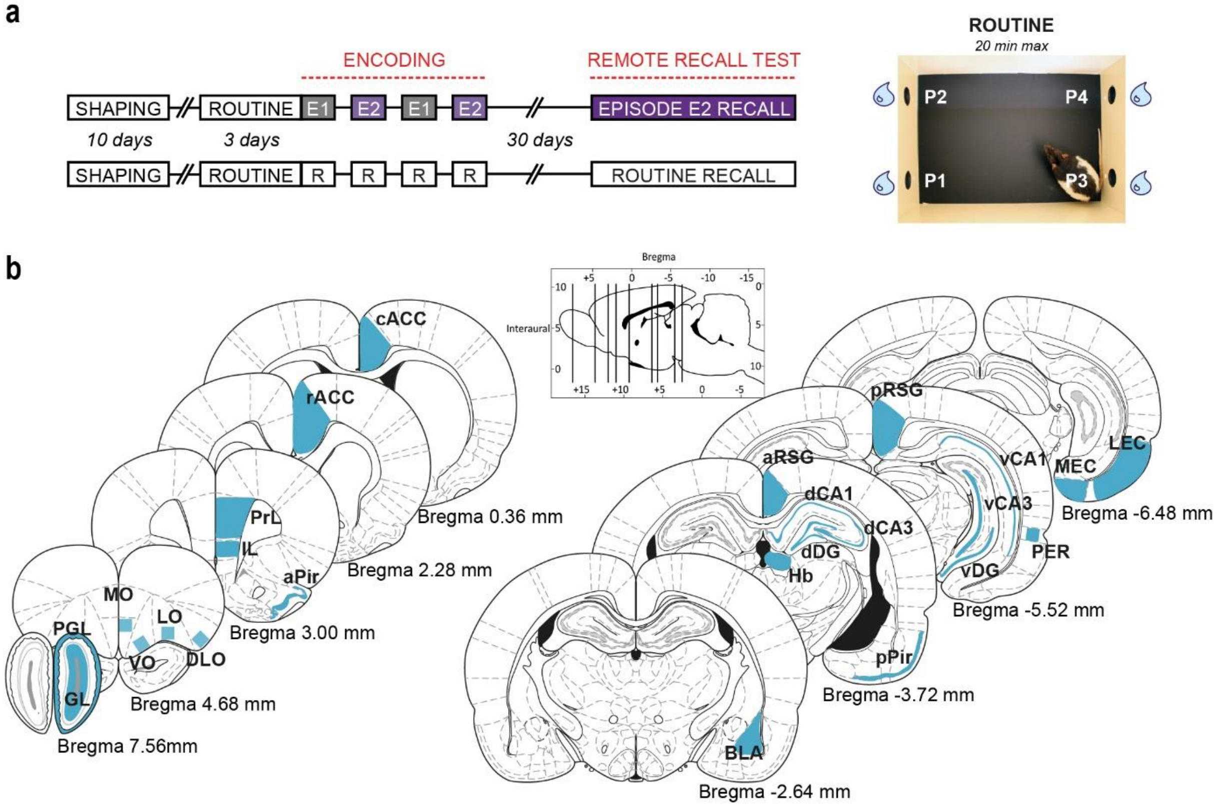


***Supplementary Figure 3. Protocol for analysing brain networks of remote episodic memories according to IEG expression***

**(a)** Timeline of the behavioural protocol for rats that experienced the episodes (n = 15) and control rats (n = 6) that experienced only routine sessions (no odour, neutral context and only water). **(b)** Rat brain coronal sections showing regions of interest selected for counting c-Fos$^+$ and Zif268$^+$ cells. The distance from the Bregma is indicated for each coronal section. The densities of IEG$^+$ cells were analysed in 23 different brain areas in 2 sections spaced by 168 µm. For the olfactory bulb (OB), counts were performed on 3 slices spaced by 504 µm. 500 µm*500 µm squares were cut for the MO, VO, LO, and DLO. GL (granular layer of the olfactory bulb); PGL (periglomerular layer of the olfactory bulb); aPir/pPir (anterior and posterior piriform cortices); MO-VO (medio-ventral), LO-DLO (dorso-lateral) part of the orbitofrontal cortex; PrL (prelimbic) and IL (infralimbic) cortices; rACC, cACC (rostral and caudal anterior cingulate cortices); aRSG, pRSG (anterior and posterior retrosplenial cortices); BLA (basolateral amygdala); Hb (habenula); dDG, vDG (dorsal and ventral dentate gyrus); dCA1, dCA3 (dorsal) and vCA1, vCA3 (ventral) hippocampus; PER (perirhinal cortex); LEC, MEC (lateral and medial entorhinal cortices).

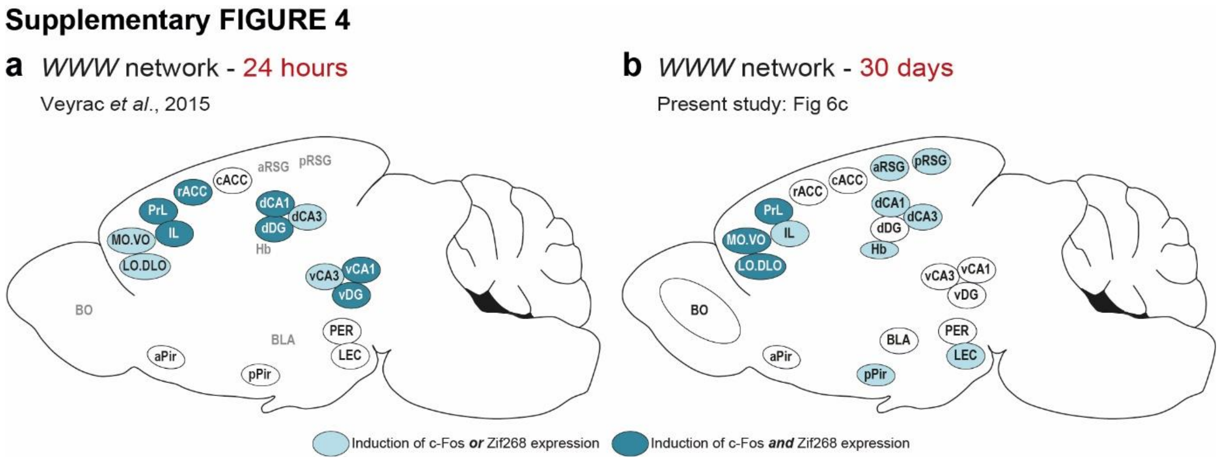


***Supplementary Figure 4. How brain networks associated with complete episodic memory evolve over time***

**(a)** Summary diagram representing brain areas recruited for *WWW* rats during recall 1 day[30] or **(b)** 30 days after episode encoding. While recent episodic memories recruited the entire hippocampus[30], only the dorsal part was recruited after a long delay. The cortical memory trace reorganized over time since the rACC was no longer in the remote memory network, whereas the OFC remains critical. The pPir and LEC, which are implicated in odour processing, were recruited only after 30 days of retention, suggesting the key role of odours in remote episodic memories. OB (olfactory bulb); aPir/pPir (anterior and posterior piriform cortices); MO.VO (medio-ventral), LO.DLO (dorso-lateral) part of the orbitofrontal cortex; PrL (prelimbic) and IL (infralimbic) cortices; rACC, cACC (rostral and caudal anterior cingulate cortices); aRSG, pRSG (anterior and posterior retrosplenial cortices); BLA (basolateral amygdala); Hb (habenula); dDG, vDG (dorsal and ventral dentate gyrus; dCA1, dCA3 (dorsal) and vCA1, vCA3 (ventral) hippocampus; PER (perirhinal cortex); LEC, MEC (lateral and medial entorhinal cortices).

#### Supplementary TABLE 1

| | | Clustering Coefficient | | Global Efficiency | |
|---|---|---|---|---|---|
| | | Data | Bootstrap p value | Data | Bootstrap p value |
| *c-Fos* | *WWW* | 0.02 | **0.008** | 0.36 | **0.050** |
| | *Where* | 0.04 | **0.010** | 0.41 | 0.723 |
| *Zif268* | *WWW* | 0.38 | **0.010** | 0.38 | **0.010** |
| | *Where* | 0.02 | **0.020** | 0.24 | 0.307 |

***Supplementary Table 1. Statistical properties of remote episodic memory brain networks.*** *Clustering coefficient* and *global efficiency* for each memory profile (*Where* and *WWW*) for c-Fos and Zif268 functional networks ($n_{WWW}$ = 9; $n_{Where}$ = 6). *P* values from bootstrap analyses show that *clustering coefficients* were significantly different from those of a random network for all graphs. Recall of incomplete episodic memories (*Where* profile) was related to less efficient functional networks since the *global efficiency* of c-Fos and Zif268 functional networks did not differ from that of a random network.

#### Supplementary TABLE 2

| a | *Where c-Fos network* | | | *WWW c-Fos network* | | |
|---|---|---|---|---|---|---|
| | **Degree** | **Strength** | **Betweenness** | **Degree** | **Strength** | **Betweenness** |
| CG | - | - | - | 4 | **6170*** | 29 |
| PGL | - | - | - | **5**** | **7530**** | 129 |
| aPir | 5 | 5541 | 7 | 3 | 4465 | **162*** |
| pPir | 5 | 5542 | 7 | 2 | 2799 | 34 |
| MO.VO | 4 | 7198 | 45 | 3 | 4931 | 34 |
| LO.DLO | 4 | 7113 | 57 | 1 | 1333 | 0 |
| PrL | 1 | -1885 | 0 | 2 | 2965 | 130 |
| IL | 3 | 1885 | 52 | 2 | 2899 | **160*** |
| rACC | 3 | 5313 | 31 | 2 | 2923 | 0 |
| cACC | 1 | 1885 | 0 | 2 | 266 | 112 |
| aRSG | 3 | 1800 | 28 | 3 | 4570 | 8 |
| pRSG | 2 | 3428 | 0 | 4 | **6674*** | 48 |
| BLA | 1 | -2000 | 0 | 1 | **-1466*** | 0 |
| Hb | - | - | - | 3 | 1832 | 34 |
| dDG | 5 | **-9198**** | 7 | 1 | **-1433*** | 0 |
| dCA1 | 1 | 1885 | 0 | 1 | 1633 | 0 |
| dCA3 | **6*** | 6970 | 75 | - | - | - |
| vDG | - | - | - | - | - | - |
| vCA1 | 5 | 5198 | 7 | - | - | - |
| vCA3 | 2 | **-3656**** | 28 | - | - | - |
| PER | - | - | - | 2 | 2932 | 64 |
| LEC | 5 | 5542 | 7 | 2 | 133 | 90 |
| MEC | 4 | **7085*** | 27 | 3 | 1366 | **166**** |
| Mean | 3.3 | 2758.1 | 21.0 | 2.4 | 2764.3 | 63.2 |

| b | *Where Zif268 network* | | | *WWW Zif268 network* | | |
|---|---|---|---|---|---|---|
| | **Degree** | **Strength** | **Betweenness** | **Degree** | **Strength** | **Betweenness** |
| CG | 1 | 1771 | 0 | - | - | - |
| PGL | - | - | - | 1 | 1523 | 0 |
| aPir | **4*** | **7198*** | 10 | 1 | 1500 | 0 |
| pPir | 3 | 5313 | 3 | 5 | 7621 | 10 |
| MO.VO | 2 | 3656 | 10 | 1 | 1366 | 0 |
| LO.DLO | - | - | - | 2 | 3252 | 0 |
| PrL | - | - | - | 1 | 1366 | 0 |
| IL | 2 | 3542 | 0 | 1 | 1500 | 0 |
| rACC | - | - | - | 5 | 7993 | 20 |
| cACC | 1 | 1885 | 0 | - | - | - |
| aRSG | - | - | - | **7*** | **11235*** | **55*** |
| pRSG | - | - | - | **8*** | **12406*** | **73**** |
| BLA | - | - | - | 1 | **-1466*** | 0 |
| Hb | - | - | - | 1 | **-1466*** | 0 |

| | | | | | | |
|---|---|---|---|---|---|---|
| dDG | 1 | 1657 | 0 | 2 | 3032 | 0 |
| dCA1 | 2 | 3428 | 2 | 5 | 8012 | 21 |
| dCA3 | 3 | 5085 | 16 | 1 | 1523 | 0 |
| vDG | - | - | - | 4 | 6574 | 1 |
| vCA1 | - | - | - | **6*** | **9159*** | 31 |
| vCA3 | 2 | 3542 | 2 | 4 | 6303 | 0 |
| PER | 3 | 5427 | 1 | 6 | **9351*** | 14 |
| LEC | 1 | 1885 | 0 | - | - | - |
| MEC | 1 | 1657 | 0 | **6*** | **9330*** | 39 |
| Mean | 2 | 3542 | 3.4 | 3.4 | 5005.7 | 13.2 |

***Supplementary Table 2. Statistical properties of functional networks of remote episodic memories***

The *degree*, *strength* and *betweenness centrality* of each node for c-Fos **(a)** and Zif268 **(b)** functional networks obtained through graph-theoretical analyses. Data are expressed as the mean values for each parameter for *Where* and *WWW* rats ($n_{WWW}$ = 9; $n_{Where}$ = 6). A bootstrap analysis determined the major hubs in each network by testing whether the node properties differed from chance (highlighted in bold red font). *$p_{bootstrap}$<0.05; **$p_{bootstrap}$<0.01. GL (granular layer of the olfactory bulb); PGL (periglomerular layer of the olfactory bulb); aPir/pPir (anterior and posterior piriform cortices); MO.VO (medio-ventral), LO.DLO (dorso-lateral) part of the orbitofrontal cortex; PrL (prelimbic) and IL (infralimbic) cortices; rACC, cACC (rostral and caudal anterior cingulate cortices); aRSG, pRSG (anterior and posterior retrosplenial cortices); BLA (basolateral amygdala); Hb (habenula); dDG, vDG (dorsal and ventral dentate gyrus; dCA1, dCA3 (dorsal) and vCA1, vCA3 (ventral) hippocampus; PER (perirhinal cortex); LEC, MEC (lateral and medial entorhinal cortices).

**Supplementary FIGURE 5**

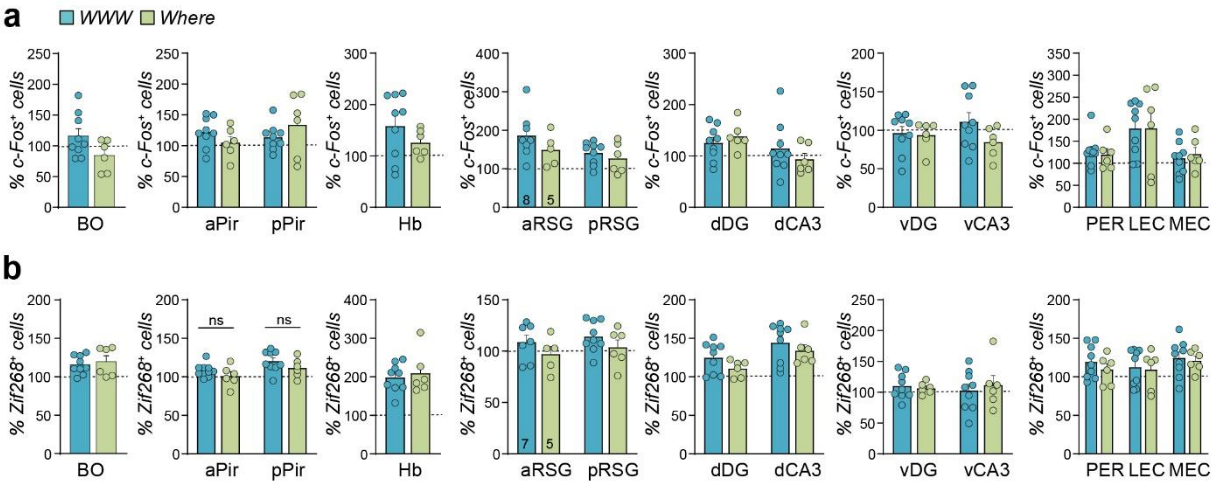


***Supplementary Figure 5. Brain areas not involved in complete remote episodic memory***

Density of c-Fos$^+$ **(a)** or Zif268$^+$ **(b)** cells in brain areas not significantly recruited in *WWW* versus *Where* rats ($n_{WWW}$ = 9; $n_{Where}$ = 6). Data were normalized to the control group (value of 100%). Mann–Whitney U tests were performed for statistical comparisons between profiles that were not significant. OB (olfactory bulb); aPir/pPir (anterior and posterior piriform cortices); Hb (habenula); aRSG, pRSG (anterior and posterior retrosplenial cortices); dDG, vDG (dorsal and ventral dentate gyrus); dCA3 (dorsal) and vCA3 (ventral) hippocampus; PER (perirhinal cortex); LEC, MEC (lateral and medial entorhinal cortices).

Supplementary FIGURE 6

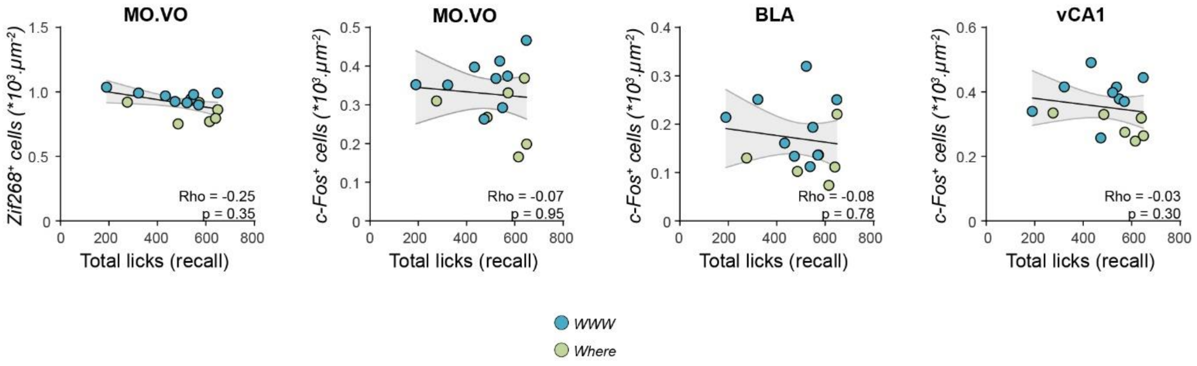


***Supplementary Figure 6. The influence of the emotional network on complete remote episodic memory is not related to the reinforcement level during the recall test***

Nonsignificant Spearman correlations between IEG (Zif268 or c-Fos) expression in emotional brain areas associated with remote memory precision in *WWW* and *Where* rats ($n_{WWW} = 9$; $n_{Where} = 6$) and the total amount of drinking during the recall test (Total licks). MO.VO (medio-ventral) of the orbitofrontal cortex; BLA (basolateral amygdala); vCA1 (ventral hippocampus).

**Supplementary FIGURE 7**

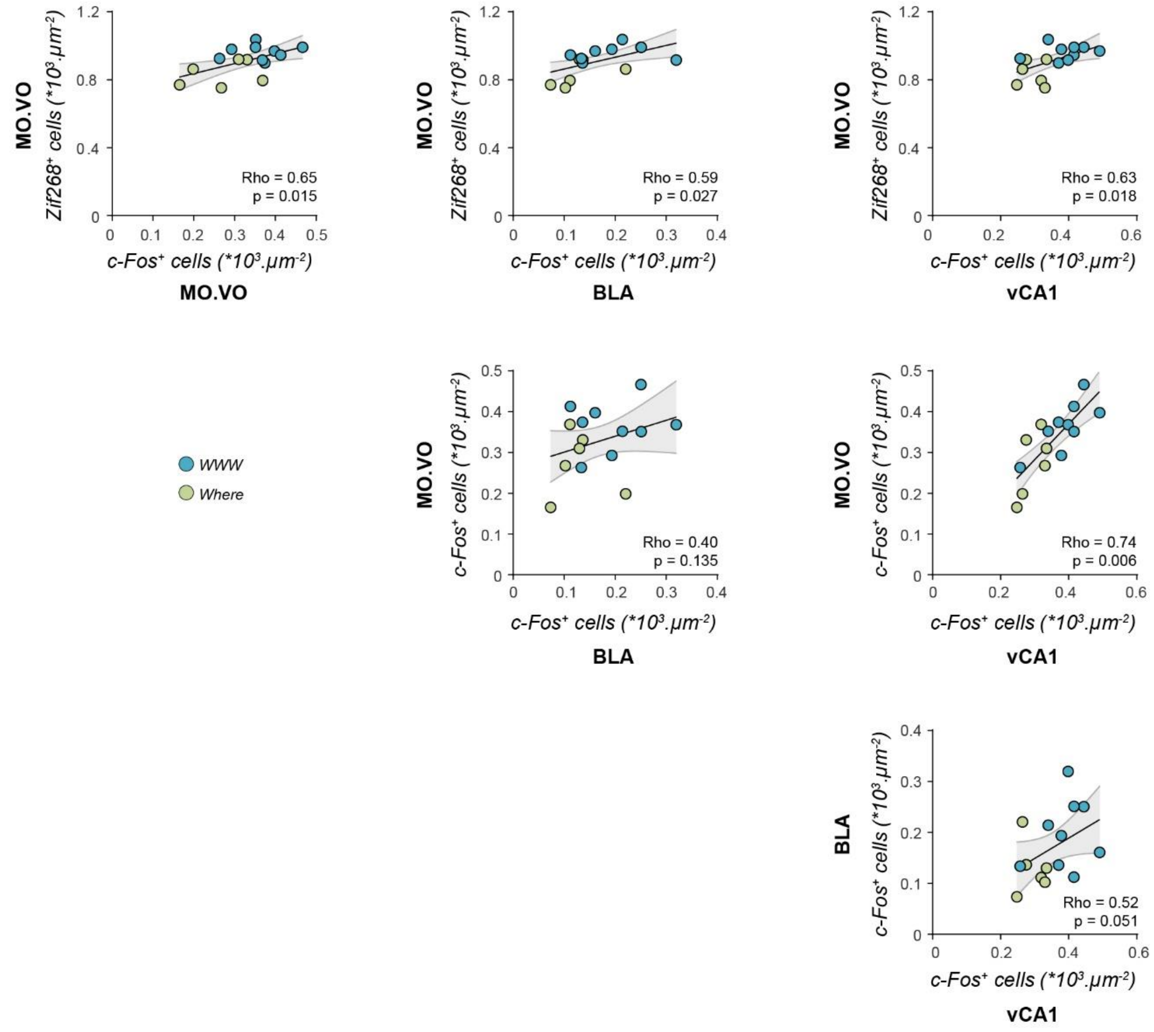


***Supplementary Figure 7. Brain areas in the emotional network involved in accurate episodic recollection are coactivated during the recall test***

Spearman correlations between c-Fos and Zif268 expression in emotional brain areas associated with remote memory precision in *WWW* and *Where* rats ($n_{WWW}$ = 9; $n_{Where}$ = 6). The coexpression of c-Fos and Zif268 was positively correlated between MO.VO, vCA1 and BLA, except for c-Fos expression in the MO.VO and BLA. MO.VO (medio-ventral) of the orbitofrontal cortex; BLA (basolateral amygdala); vCA1 (ventral hippocampus).

**Supplementary TABLE 3**

| Mediation of… | through… | Mediate effect | | Direct effect | |
|---|---|---|---|---|---|
| …for predicting memory precision | | % of mediation | p value | % of direct effect | p value |
| **Zif268 MO.VO** | c-Fos BLA | 30 | 0.08 | 70 | **0.02** |
| **Zif268 MO.VO** | c-Fos vCA1 | 9 | 0.58 | 91 | **0.01** |
| **Zif268 MO.VO** | c-Fos MO.VO | 1 | 0.95 | 99 | **0.01** |
| c-Fos BLA | **Zif268 MO.VO** | 43 | **0.02** | 57 | 0.09 |
| c-Fos vCA1 | **Zif268 MO.VO** | 76 | **0.02** | 24 | 0.58 |
| c-Fos MO.VO | **Zif268 MO.VO** | 100 | **0.01** | 0 | 0.99 |

***Supplementary Table 3. Statistical properties of mediation analyses on the emotional network involved in accurate episodic recollection***

Values and statistical data of mediation analyses that determined relationships between brain areas recruited and remote episodic memory accuracy (n = 15). MO.VO (medio-ventral) of the orbitofrontal cortex; BLA (basolateral amygdala); vCA1 (ventral hippocampus).
